# Dynamic regulation of engineered T7 RNA polymerases by endogenous metabolites

**DOI:** 10.1101/2024.08.07.607023

**Authors:** Zachary T. Baumer, Matilda Newton, Lina Löfstrand, Genesis Nicole Carpio Paucar, Natalie G. Farny, Timothy A. Whitehead

## Abstract

For many enzymes, activity is regulated post-translationally by endogenous metabolites. Designing liganded control of essential activities like transcription would advance predictive control of biological processes, a fundamental goal of synthetic biology. Here we demonstrate that full-length, single subunit T7-derived RNA polymerases (T7 RNAP) can be controlled by physiologically relevant concentrations of indoles. We used rational design and directed evolution to identify T7 RNAP variants with minimal transcriptional activity in the absence of indole, and a 29-fold increase in activity with an EC_50_ of 344 *µ*M. Indoles control T7-dependent gene expression exogenously, endogenously, and inter-cellularly. We also demonstrate indole-dependent bacteriophage viability and propagation in *trans*. Specificity of different indoles, T7 promoter specificities, and portability to different bacteria are shown. Our <u>l</u>igand <u>a</u>ctivated <u>R</u>NA <u>p</u>olymerases (LARPs) represent a new chemically inducible platform immediately deployable for novel synthetic biology applications, including for modulation of synthetic co-cultures.

## INTRODUCTION

Following the discovery of the genetic code, “the first secret of life”, Monod described the “second secret of life” - that biological macromolecules and small molecules interact in complex ways to rapidly control metabolic fluxes in response to ever changing environmental conditions^1^. Deciphering these secrets have culminated in our ability to engineer control over many aspects of cellular biology ^2–4^. Although natural metabolite-responsive enzymes are ubiquitous in cellular life ^5–7^, such enzymes have proven difficult to engineer or design. Most examples contain a synthetic signature with complicated, abiological, protein fusions and repurposing of ligand binding domains ^8–12^. We chose to design dynamic metabolite control over the general transfer of information of DNA to RNA^13^ using T7 RNAP. T7 RNAP and derivatives have been studied extensively ^14–16^, are used to synthesize mRNA for medical applications ^17,18^, and are the dominant transcriptional control mechanism employed in recombinant protein expression in bacteria ^19^. However, unregulated *in vivo* expression of T7 RNAP is usually toxic owing to its high basal activity ^20,21^. We chose indoles as a metabolite class because they are inexpensive, used for interspecies communication for biofilm formation ^22^ and in the gut microbiome, and are associated with various disease states ^23–25^. **L**igand **a**ctivated **R**NA **P**olymerases (LARPs) with minimal activity in the absence of indoles would allow user-defined transcriptional processes in diverse bacteria with little toxicity to the host.

Herein we use rational design to engineer the full-length, single subunit T7 RNA Polymerase to be controlled by physiologically relevant concentrations of indole. We optimize our LARPs through directed evolution to yield LARP-I with minimal transcriptional activity in the absence of indole, and a 29-fold increase in activity with an EC_50_ of 344 *µ*M. We utilize LARP-I in several contexts to show that indole control T7-dependent gene expression exogenously, endogenously, and inter-cellularly. We also demonstrate indole-dependent bacteriophage viability and propagation in *trans*. Specificity of different indoles, T7 promoter specificities, and portability to different bacteria are shown. Our <u>l</u>igand <u>a</u>ctivated <u>R</u>NA <u>p</u>olymerases (LARPs) represent a new chemically inducible platform immediately deployable for novel synthetic biology applications, including for modulation of synthetic co-cultures.

## RESULTS

### Design, engineering, and optimization of LARPs

We used a chemical recovery of structure approach to identify initial LARPs ^26,27^ (**Fig 1A**). We performed glycine scanning mutagenesis of 15 Trps buried in the transcription initiation complex of T7 RNAP ^14^, expressed proteins as N-terminal His_6_ tag fusions, purified proteins over nickel columns, and assessed these proteins for indole-dependent activity using a modified *in vitro* transcriptional assay ^28^. In this assay, potential LARPs are incubated with linear dsDNA containing a T7 promoter sequence driving expression of an RNA Spinach or Pepper aptamer ^29,30^. The amount of transcript is proportional to fluorescence, and the rate of fluorescence is determined in the presence and absence of 1 mM indole (**Fig S1**). Six of the 15 designs expressed in the soluble fraction of *E. coli* lysates (**Fig S2**), and none showed indole responsiveness relative to the solvent control. Two different computational methods ^31,32^ predicted that the glycine mutations for the insoluble designs were highly destabilizing (**Fig S3**). We hypothesized that expression of the insoluble variants could be rescued in a thermally stabilized (TS) background. Upon transfer to the TS background^33^, two variants were rescued and responsive at 1 mM indole (**Fig S4)**. The best design, LARP_V1_ (T7 RNAP S430P, N433T, S633P, W727G, F849I, F880Y), showed a 3.2-fold increase in transcription rate (s.d. = 0.9 [2.3,4.1]; n=6) (**Fig 1B)** with an approximate EC_50_ of 152 μM indole (95% c.i. 117-200 μM; n=6) **(Fig S5**).

**Fig. 1:**
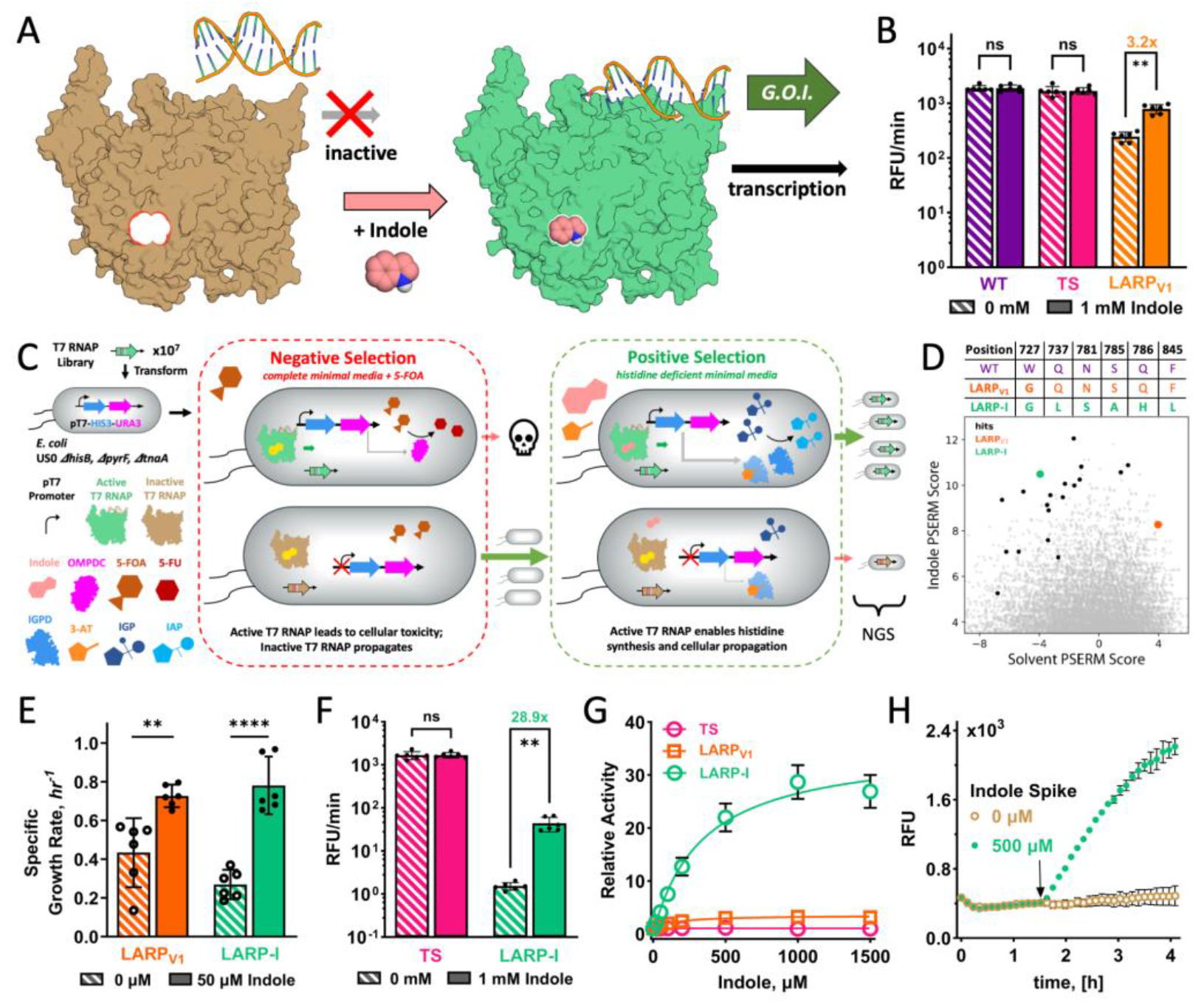
Development of a ligand-activatable RNA polymerase responsive to indole. **[A]** Chemical recovery of function approach for T7 RNAP. Mutating a buried tryptophan in T7 RNAP disrupts the ability of the RNAP to transcribe DNA to RNA. This activity is recovered in the presence of indole. **[B]** *In vitro* transcriptional assays using Peppers aptamer in the absence or presence of 1 mM indole for the indicated variants. TS_W727G_ is also known as LARP_V1_. **[C]** Schematic of the bacterial-1-hybrid selection system developed for identification of LARPs. The selection enables positive and negative selection. **[D]** PSERM scores for library variants for indole vs. selection on a solvent control. The Pareto front of high score in the presence of indole and low score for the solvent contains variants (large green circle for LARP-I, black circles for other tested variants) predicted to have improved or maintained indole responsive growth with significantly less constitutive activity compared to the starting construct (LARP_V1_, orange larger circle). The sequence profile above the plot contains the LARP-I and LARP_V1_ specific mutations relative to the wild-type T7 RNAP. **[E]** Specific growth rates of LARP constructs in the presence and absence of 50 *µ*M indole under growth in selective minimal media without histidine supplementation and with 1 mM 3-AT. **[F-G]** *in vitro* transcriptional assay using the Peppers aptamer with indicated indole concentrations for LARP-I, LARP_V1_, and TS. Panel F shows activity represented as RFU/min, and panel G shows a relative activity for each variant normalized to one in the absence of indole. **[H]** *in vitro* transcriptional activity time course. A delayed spike of 500 *µ*M indole shows rapid activation of LARP-I transcription. Statistics and p-values: **: p-value<0.01, ****: p-value<0.0001; ns Not significant (p-value >0.05). RFU/min given as mean and standard deviation (B, F), Welch ANOVA using Brown-Forsyth and corrected for multiple comparisons, Specific Growth Rate given as mean and 95% C.I. Welch’s unpaired two-tailed t-test and Tukey’s post-hoc test. Relative Activity (G) reported as mean and SEM error bars, the RFU in the delayed indole trace (H) is given as a range. All data is presented in SOURCE DATA FILE.

In the absence of indole, LARP_V1_ has 13.1% (s.d. = 3.1%; n=6) activity relative to native T7 RNAP (**Fig 1B**). Near zero basal activity is required for most relevant applications. To identify low background, highly inducible LARPs we optimized a previously described dual positive and negative bacterial selection ^34,35^ (**Fig 1C**). In this system, a plasmid construct containing a T7 promoter upstream of HIS3 and URA3 genes is transformed into an *E. coli* strain (i.) incapable of producing endogenous indole; and (ii.) auxotrophic for histidine and uracil (*E. coli* US0 ***𝜟****hisB, 𝜟pyrF, 𝜟tnaA*; **Fig S6**). Growth on minimal salts media deficient in histidine requires T7-dependent transcription of HIS3. Constitutive LARPs can be counter selected against by growth on media supplemented with 5-fluoro-orotic acid (5-FOA), as the expressed URA3 gene product will convert 5-FOA to the cytotoxic 5-fluorouracil. Extensive expression engineering of LARP_V1_ was necessary to move into the appropriate ‘Goldilocks’ zone of the selection. High LARP_V1_ expression resulted in high toxicity in the absence of selection conditions, while too low of basally expressed LARP_V1_ did not recover differential growth in the presence of indole under selection conditions. We found that expression of a LARP_V1_ construct from a low copy number plasmid (ORI p15A), replacing the ATG start codon with GTG, and addition of an N-terminal Degron tag were needed to tune the appropriate expression window ^36^ (**Fig S7**).

This method was used to select low basal activity LARPs from a library of approximately 11 million members ^37^. A combinatorial library containing mutations at nine positions within 6 Å of position 727 was constructed by a cassette-based Golden Gate protocol using synthetic DNA containing degenerate codons ^37^. Transformed cells were passed through a round of negative selection using 1 mM 5-FOA. Surviving members of this library were then selected for activity in the presence and absence of 50 *µ*M indole. Deep sequencing of the selected and input libraries^38^, followed by PSERM analysis ^39^, revealed a Pareto front of candidate indole-specific LARPs (**Fig 1D, Fig S8)**. Of the 20 variants on the Pareto front that were cloned and tested, 11 LARPs passed plate-based assays and showed statistically significant (One-way ANOVA, Tukey’s post-hoc test, p-value<0.03 for all samples) indole-specific growth rate increases in defined minimal media (**Fig S9; Table S1**). LARP-I (K378R, S430P, N433T, S633P, W727G, Q737L, N781S, S785A, Q786H, F845L, F849I, F880Y) is marked by minimal growth rate in the absence of indole, and near-maximal growth rate with 50 *µ*M indole under selection conditions **(Fig 1E)**. We expressed and purified LARP-I (**Fig S5**) to evaluate its binding properties *in vitro*. The basal activity of LARP-I is less than 0.1% of the activity of parent enzyme T7 RNAP TS, the EC_50_ for indole is 344 *µ*M (95% c.i. 193-667 *µ*M; n=6), and the dynamic range for indole responsiveness is 28.9 at 1 mM (s.d. = 12.2; n=6) **(Fig 1F-G)**. To evaluate the timescales of indole activation, we assayed the time-dependent fluorescence; a late-addition spike of indole also shows post-translational temporal control over transcription within minutes **(Fig 1H)**. Thus, LARP-I identified from our selection shows low basal activity and post-translational, indole-inducible transcription with a high dynamic range *in vitro*.

### LARPs control gene expression exogenously, endogenously, and in co-cultures

We evaluated the ability of LARP-I to control indole-dependent gene expression in a variety of contexts. A plasmid encoding a sfGFP expression marker downstream of a T7 promoter sequence was transformed into *E. coli* expressing LARP-I and lacking tryptophanase (ΔtnaA) (**Fig 2A**). LARP-I shows low constitutive expression in the absence of indole (< 1.5-fold RFU/OD_600_ above negative control), and a 17.1-fold increase in sfGFP reporter expression at 500 *µ*M indole (s.d. =3.7, n=3) **(Fig 2B)**. The maximum response under the conditions of the assay was 40.8% (s.d. = 9.0%, n=3) of the response of T7 RNAP R632S under the same promoter, plasmid, and condition (**Fig 2B**). Thus, external addition of indole is sufficient to drive high levels of gene expression.

**Fig. 2:**
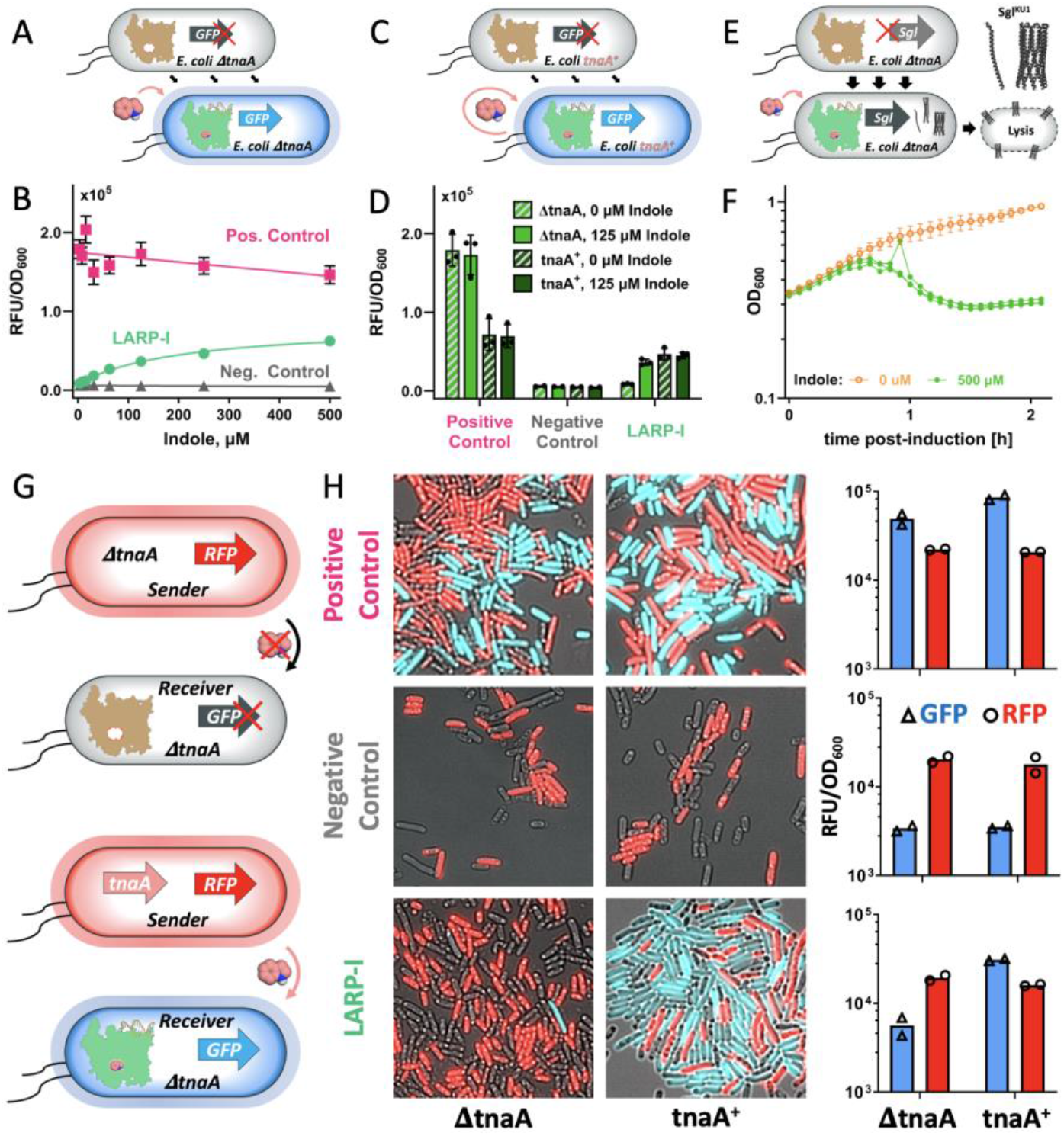
LARP-I allows indole control of gene expression exogenously, endogenously, and intercellularly. **[A]** Cartoon of predicted results of experiment with external addition of indole. Exogenous indole added to LARP-I containing *E. coli* and the tryptophanase gene knocked out enables ligand dependent gene expression measured by expression of a fluorescent reporter. **[B]** GFP RFU normalized by cell density as a function of supplemented indole concentration. Positive control represents an expression strain using T7 RNAP^R632S^, and negative control expresses the catalytic knockout T7 RNAP^R632S/Y639A^. Error bars represent 1 s.e.m., n≥3. **[C]** Cartoon of predicted results using an *E. coli* strain capable of producing indole. *E. coli* naturally expresses tryptophanase, and produces indole through tryptophan metabolism. **[D]** GFP RFU normalized by cell density for indicated strains and in the presence and absence of 125 *µ*M indole. Positive control (T7 RNAP^R632S^) and negative control (T7 RNAP^R632S/Y639A^) are the same as in panel B. Endogenous activation of LARP-I is similar to exogenous activation at 125 *µ*M indole addition in direct comparison of strains with and without the tnaA knockout. Error bars represent 1 s.d., n≥3. **[E]** Predicted effects of the expression of single gene lysin (Sgl^KU1^) under a T7 promoter with co-expression of LARP-I. Indole activates LARP-I, leading to cell lysis. **[F]** OD_600_ vs. time after induction of the culture with 500 *µ*M indole. Ethanol was used as a vehicle control for the 0 *µ*M control. Indole-dependent cell lysis of *E*.*coli* expressing Sgl^KU1^ occurs within 45 minutes. Error bars for 0 *µ*M indole represent s.e.m., n=3. The three biological replicates are plotted for the 500 *µ*M indole experiment. **[G]** Cartoon of expected activation of receiver strain by the sender strain. **[H]** Confocal microscopy of representative co-cultures; panels to the right show RFU/OD_600_ bulk population measurements for the co-cultures shown by microscopy. Positive and negative T7 RNAP controls are the same as in panels B and D. All data is presented in SOURCE DATA FILE.

To determine whether LARP-I can be activated by endogenous metabolites, we transformed plasmids containing the reporter sfGFP and LARP-I in *E. coli* expressing tryptophanase. In planktonic cultures and near stationary phase, *E. coli* with active tnaA produce mM concentrations of indole (**Fig 2C**). *E. coli* tnaA^+^ activates similar levels of gene expression as *E. coli* ΔtnaA with 125 *µ*M indole (**Fig 2D**). No statistically significant difference in gene activation occurs in the presence or absence of 125 *µ*M indole in *E. coli* tnaA^+^ (**Fig 2D;** p-value=0.7602, Welch’s two-tailed t test). *E. coli* tnaA^+^ expressing LARP-I shows a 12.4-fold increase in gene expression over the ΔtnaA in the absence of exogenous indole (s.d.=2.6, n=3) and 62.2% gene expression compared to T7 RNAP^R632S^ in direct comparison (s.d. = 21.3%, n=3).

Receiver strains could release a biological payload in response to indole either through cell lysis or secretion. To demonstrate that indole could be used to induce lysis, we screened a panel of previously described single gene lysins ^40,41^ and identified several that lysed *E. coli* under induction with an arabinose-inducible promoter, including Sgl^KU1^. Indole induction of Sgl^KU1^ by LARP-I expressing *E. coli MG1655 Δ*tnaA resulted in cell lysis within 45 minutes (**Fig 2E. Fig S11**). Therefore, LARP-I can be controlled by physiologically relevant concentrations of an endogenous metabolite. Additionally, LARP-I enables indole-mediated delivery of intracellular cargo through cell lysis.

Microbiome engineering would benefit from additional bottoms-up quorum signaling circuits ^42–44^. To determine whether LARP-I can be used for indole-dependent intercellular signaling, we constructed a sender strain (*E. coli* US0 ***𝜟****hisB, 𝜟pyrF, RFP*^*+*^) that produces indole. As a control, we also prepared a sender strain deficient in indole production (*E. coli* US0 ***𝜟****hisB, 𝜟pyrF 𝜟tnaA, RFP*^*+*^). For the receiver strain, we constructed (*E. coli* US0 ***𝜟****hisB, 𝜟pyrF 𝜟tnaA)* with the LARP-I/sfGFP reporting system (**Fig 2E**). Additional receiver strains encoding T7 RNAP^R632S^ and a catalytically inactive T7 RNAP R632S/Y639A were included as positive and negative controls, respectively. Population measurements of fluorescence show a 12.9-fold difference in sfGFP expression (n=2; [7.1,19.3]) for the LARP-I receiver strain when co-cultured with the sender strains able or unable to produce indole. No differences in sfGFP expression were observed for the negative control receiver strains regardless of the sender strain (**Fig 2F**). Fluorescence microscopy of the co-cultures shows that most of the individual LARP-I receiver cells are activated in a co-culture with the indole-producing sender strain (representative data shown in **Fig 2F)**, while few are activated by the indole-deficient sender strain. Thus, LARP-I can be part of novel receiver circuits that allows intercellular communication between indole sender strains and LARP-I containing receiver strains.

### LARPs enable ligand-dependent bacteriophage viability *in trans*

To test whether LARP-I enables indole dependent phage propagation *in trans*, we infected *E. coli* expressing LARP-I with T7 **Δ**gp1 bacteriophage that does not contain the vital gp1 gene (T7 RNAP) ^45^ (**Fig 3A**). In LARP-I expressing strains, a 10^4^ increase in countable plaques was observed between 0 and 500 μM indole (**Fig 3B**), and the number of plaques at 500 μM was indistinguishable between strains expressing WT T7 RNAP and LARP-I (**Fig 3B**). To evaluate whether phage infection efficiency is ligand-dependent, we tested LARP-I expressing strains using a phage infection kill-curve assay ^46^. At increasing indole concentrations, a more robust phage infection (time to clearance) was observed **(Fig 3C)**. Thus, indole dependent infection and propagation enabled by LARP-I allows interkingdom communication between phage and bacteria.

**Fig. 3:**
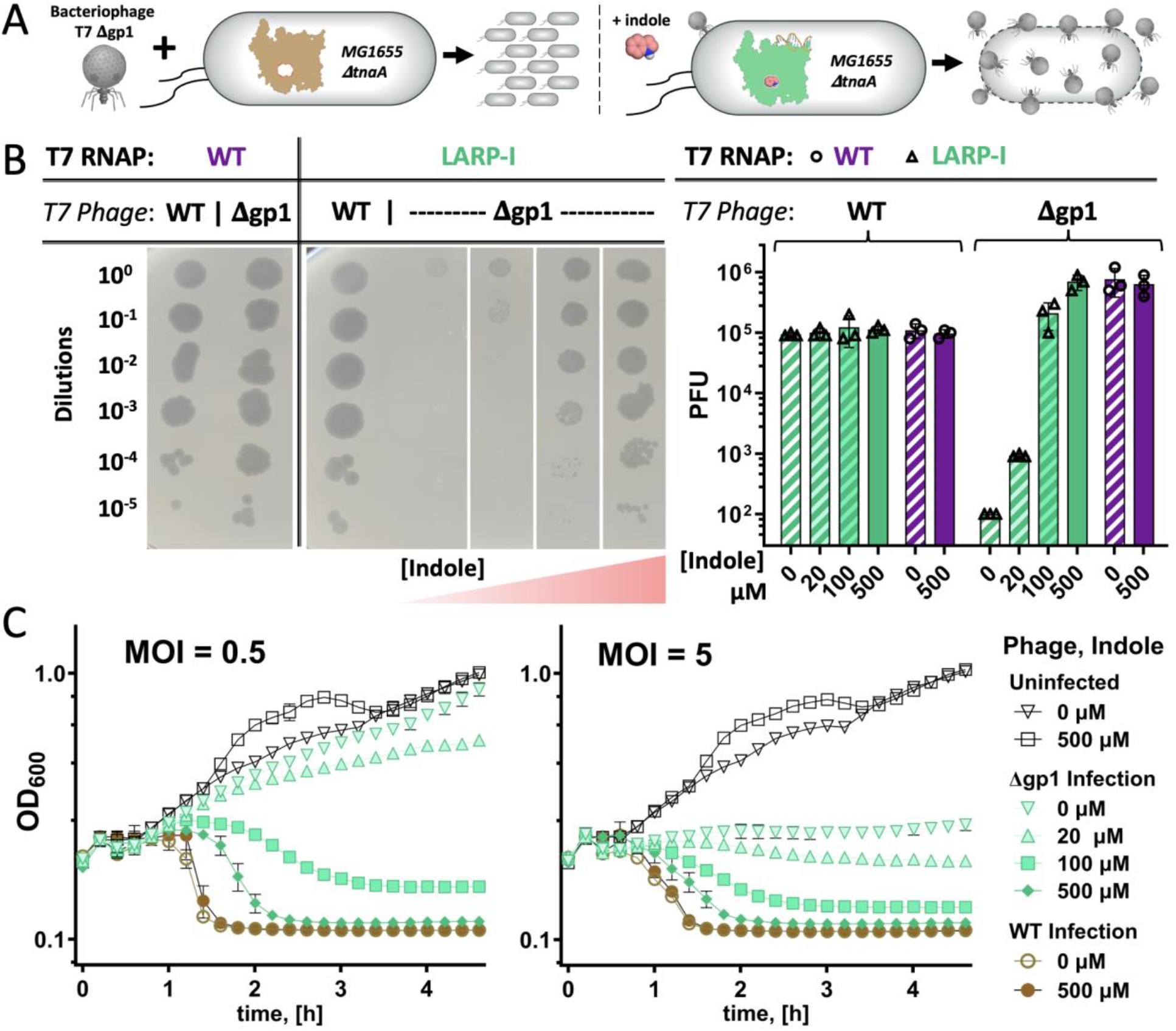
LARP-I controls T7 bacteriophage viability and propagation in trans. **[A]** Cartoon of phage experiment with the phage RNAP supplied in trans. Bacteriophage T7 Δgp1 (ΔT7 RNAP) requires active T7 RNAP to propagate. Bacteria containing LARP-I do not propagate phage in the absence of indole, but yield robust phage infection in the presence of indole. **[B]** Phage plaque formation, and quantification as plaque forming units (PFU), for different T7 phage, T7 RNAPs, and indole concentrations. **[C]** OD_600_ vs. time after infection for indicated combinations of phage and indole for LARP-I expressing *E. coli* MG1655 ΔtnaA. Two different multiplicities of infection (MOI) are shown. Phage infection kinetics are indole-dependent in kill-curve assays approaching WT phage infection rates at 500 ⌈M indole. Error bars represent 1 s.d.,(n=3). All data is presented in SOURCE DATA FILE.

### LARPs are portable for different bacteria, DNA promoter specificities, and ligands

New synthetic biology applications would be enabled if LARPs could be demonstrated to function in different organisms, with different promoter sequence specificities, and with different controlling ligands. To determine the portability of LARP-I to other organisms, we integrated a sfGFP reporter driven by a canonical T7 promoter and constitutively expressed LARP-I in *Pseudomonas putida* AG4775 ^47^ (**Fig 4A**). *P. putida* AG4775 does not endogenously express indole (**Fig S11A**) and growth is retarded somewhat by sub-mM concentrations of indole (**Fig S11B**). Nevertheless, LARP-I shows robust activation of a GFP reporter at increasing indole concentrations, as measured by flow cytometry (**Fig 4B, S11C**).

**Fig. 4:**
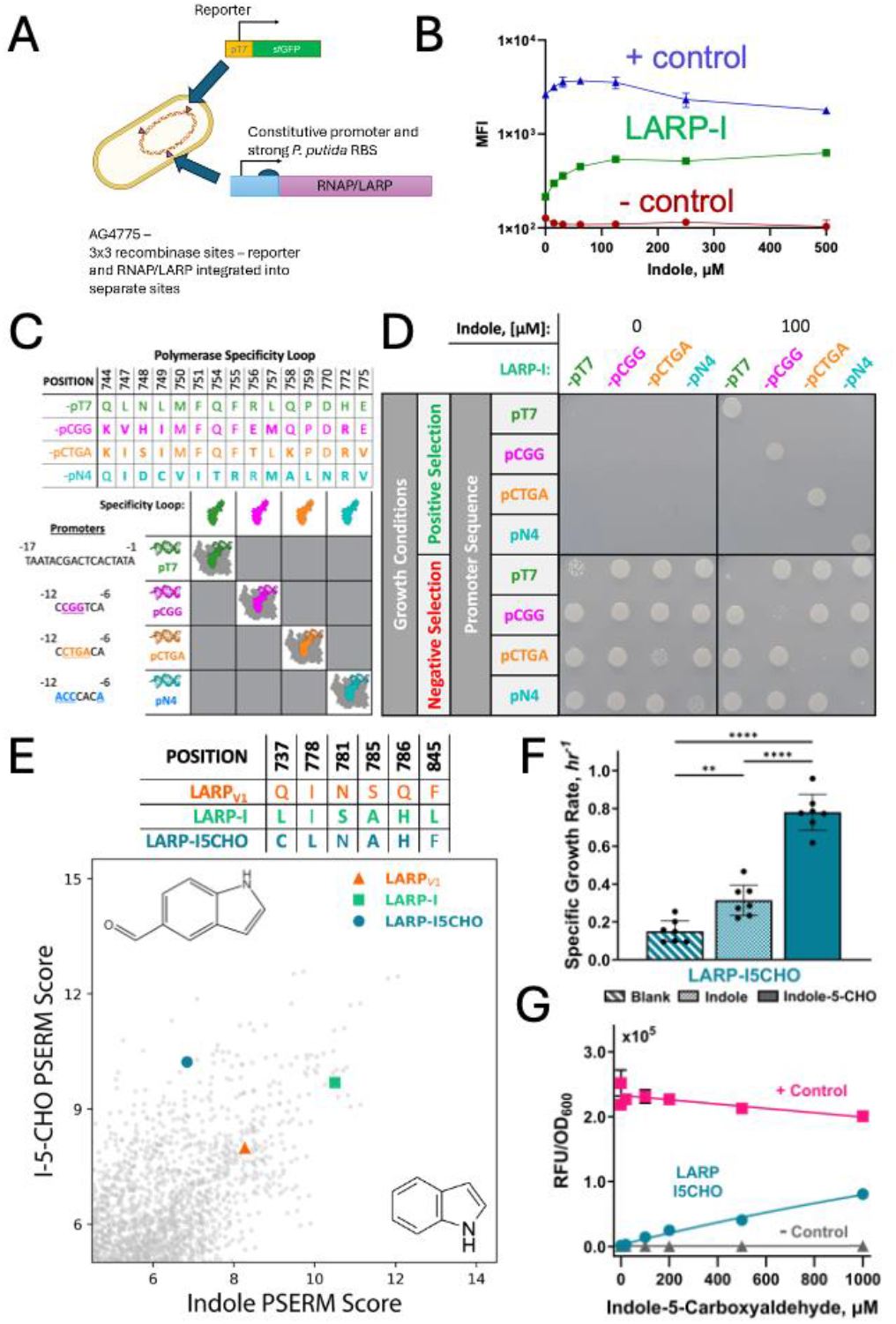
The versatility and orthogonality of LARPs are demonstrated for different bacteria, different promoter sequence specificities, and for different controlling ligands. **A**. Cartoon of experimental construction of *P. putida* LARP constructs. **B**. GFP Mean fluorescence intensity (MFI) as a function of indole for LARP-I, positive control (T7 RNAP^R632S^), and negative control (T7 RNAP^R632S/Y639A^). Error bars represent 1 s.e.m. of n=3. **C**. Hybrid LARPs with engineered polymerase specificity loops. Sequence variation at the specificity loops for LARP-I-pCGG, LARP-I-pCTGA, and LARP-I-pN4 are shown relative to the original LARP-I which has activity on the canonical pT7 promoter sequence. **D**. Selection of *E. coli* strains in the presence and absence of 100 *µ*M indole. Strains contain the indicated hybrid polymerases and promoters driving URA3 and HIS3 expression. Positive selection conditions are growth on media lacking histidine supplemented with 1 mM 3-AT, while negative selection uses complete growth media containing 1 mM 5-FOA. **E**. PSERM scores of library variants from selection on indole-5-carboxyaldehyde (I-5-CHO) vs. indole. Larger symbols represent indicated variants. Sequence differences between variants are shown. **F**. Specific growth rate on media lacking histidine and supplemented with 1 mM 3-AT. Indole and I-5-CHO are included at 50 *µ*M. Statistics and p-values : **: p-value<0.01, ****: p-value<0.0001. **G**. E. coli USO ΔtnaA with an sfGFP reporter plasmid driven by pT7 and different T7 RNAPs (positive control: T7 RNAP^R632S^; negative control: T7 RNAP^R632S/Y639A^). GFP RFU normalized to OD_600_ as a function of I-5-CHO concentration for the indicated strains at 22 hours post induction. Error bars represent 1 s.d. (n=3). All data is presented in SOURCE DATA FILE.

Previous studies engineered T7 RNAPs that respond selectively to different promoter sequences^20,36,48^. These engineered T7 RNAPs all contained mutations in their specificity loops (positions 739-766) adjacent to the LARP-I mutations. To determine the portability of the LARP-I mutations, we engineered hybrid polymerases containing both the LARP-I and the specificity loop mutations (**Fig 4C**). At the same time, we also constructed plasmids containing alternative promoters (pCGG, pCTGA, pN4) driving URA3 and HIS3. If the hybrid polymerases were functional, positive and negative selection using the auxotrophic strain *E. coli* US0 ***𝜟****hisB, 𝜟pyrF, 𝜟tnaA* would result in the same growth phenotype as with pT7 and LARP-I. If the hybrid polymerases were specific, then for non-cognate pairs positive selection would result in no growth while negative selection would result in growth. To test this hypothesis, we transformed *E. coli* with the combinatorial set of hybrid polymerases and promoters. For the hybrid polymerases tested, indole-dependent growth was observed only for cognate polymerase-promoter pairs (**Fig 4D**). Consistent with this, ligand-dependent toxicity was observed only for those same cognate polymerase-promoter pairs (**Fig 4D**). Thus, the LARP-I mutations are transferable to engineered T7 RNAPs with altered promoter specificity.

To determine whether LARPs could be engineered for other controlling ligands, we repeated the selection and analysis shown in **Fig 1C** for the indole derivative indole-5-carboxyaldehyde (I-5CHO; **Fig S8**). Following PSERM analysis comparing I-5CHO vs. indole scores (**Fig 4E**) we identified LARP-I5CHO (K378R, S430P, N433T, S633P, W727G, Q737C, I778L, S785A, Q786H, F845L, F849I, F880Y). As determined by growth on media without histidine, *E. coli* US0 ***𝜟****hisB, 𝜟pyrF, 𝜟tnaA* expressing LARP-I5CHO grows minimally in the absence of I-5CHO and is selectively activated by I-5CHO over indole (**Fig 4F**). Expression of LARP-I5CHO in the same reporter strain shown in **Fig 2A** revealed I-5CHO dependent sfGFP expression, with the overall activation on the same order of magnitude as constitutive expression of T7 RNAP^R632S^(**Fig 4G**). Thus, LARPs can be engineered to be specific to other indoles.

The LARPs demonstrated here have minimal basal activity and large indole-dependent increases in activity. The LARP-specific mutations centered around W727 are located beneath the interface with the allosteric inhibitor T7 lysozyme ^49^. T7 lysozyme traps T7 RNAP in the initiation complex where, while it is able to bind DNA, only produces abortive transcripts. It is possible that a similar allosteric mechanism occurs for the LARPs. The binding of LARPs to pT7 DNA was tested by fluorescently labeling one strand of the DNA duplex and performing fluorescence anisotropy using titration of T7 RNAP ^50,51^. TS recognized pT7 with a K_D_ of 1.8 nM at 25 and 37°C [1.41-2.22] and in the presence or absence of indole, consistent with previous literature values of WT ^50^ (**Fig S12)**. In the absence of indole LARP-I also bound pT7 DNA, albeit with a reduced K_D_ of 26 nM [24.17-26.94] (**Fig S12)**. In the presence of indole the affinity for pT7 DNA modestly improved to 14.6 nM [11.92-17.59] (**Fig S12)**. We also used circular dichroism to evaluate the secondary structure and the apparent melting temperature of LARP-I with and without indole. There was no significant difference in the secondary structure of LARP-I relative to the TS background (**Fig S13A**). While T7 RNAP is largely an alpha helical protein, the locations of the LARP mutations are close to beta strands from positions 720-770. There were no differences in ellipticity as a function of temperature at 222 nm (measuring alpha helical content) in the presence or absence of indole (**Fig S13B**). On the other hand, the presence of indole stabilized other secondary structures (measured by 208.5 nm ellipticity) for LARP-I (**Fig S13C)**, consistent with the expected binding location. This stabilization is also congruent with modest differences for the affinity of LARP-I with pT7 DNA observed in the presence of indole. In the absence of indole, LARP-I is able to recognize pT7 DNA. While these initial results support a similar mechanism of activation as T7 lysozyme binding inhibition, further biophysical or biochemical experiments are needed before exact mechanisms are established.

## DISCUSSION

The LARPs developed here represent a new chemically inducible system that should function in diverse bacteria with minimal modification. This system supplies an urgent need for the synthetic biology community of predictive gene expression control with an alternative controlling ligand. Further, LARPs can sense and respond to the native metabolite indole, which allows LARP receiver strains to be constructed for bottoms-up manipulations of microbial communities. We also demonstrated that LARPs can respond selectively to an indole derivative. LARPs may find applications in inducible or dynamic control of gene expression in bioreactors, for metabolite control of engineered phage therapies, or to perform user-defined operations in the gut microbiome or other mixed microbial communities producing indole and indole derivatives.

More fundamentally, these results show that dynamic regulation of protein activity by ligands can emerge given sufficient sampling of different evolutionary trajectories. Ligand-responsive enzymes ^5^ are ubiquitous in nature, particularly for central metabolism where precise control of fluxes are paramount. Completely distinct mechanisms of regulation in homologous proteins suggest that such dynamic regulation is a natural consequence of protein and ligand co-localization, can emerge by differing evolutionary trajectories, and that such emergence should be a relatively common evolutionary event ^52^. Under the hypothesis that ligand-responsive enzymes arise spontaneously from the laws of mass action and colocalization, our design and directed evolution strategy in this work could be generalized and extended to other high value enzymes for which agonist switching activity would be strongly desired. Given that our LARP mutations are localized to a known allosteric network for T7 RNAP, development of new ligand-responsive enzymes may be improved with sufficient prediction of allosteric locations in known enzymes ^53^.

## Acknowledgements

We would like to acknowledge J. Yesselman for the gift of HBC530, J.J. Bull and I. Molineux for the gift of Δgp1 T7 bacteriophage; B. Seelig for the gift of an T7 RNAP expression construct; A. Whiteley, E. Kibby, and U. Tak for help with phage; A. Hren, A. Avramov, C. Huffine, N. Skillin & J. Cameron for help with imaging; M.D. Smith for help with PSERM implementation; S. Kennedy, A. Erbse, H. Steiner, and R. Garcea for help with biophysical assay development. We also thank the Shared Instruments Pool (RRID: SCR_018986) of the Department of Biochemistry at the University of Colorado Boulder for the use of the CD spectrometer and the Tecan Spark plater reader. The CD is funded by NIH Shared Instrumentation Grant **S10RR028036**.

## Author contributions

Z.T.B. and T.A.W. developed the idea; Z.T.B., M.N., G.N.C.P., N.F., and T.A.W. designed experiments; Z.T.B., M.N., L.L., G.N.C.P. performed experiments; Z.T.B., G.N.C.P. performed data analysis; Z.T.B. and T.A.W. wrote the manuscript with contributions from all co-authors.

## Data and materials availability

The raw sequencing files are publicly available as SRA depositions under the following project numbers (BioProject ID: PRJNA1136062; **<u>Numbers deposited upon publication</u>**). Raw image files for plate growth assays, plaque assays, and fluorescence microscopy are available at https://zenodo.org/records/12676214. All other data are available in the manuscript or the supplementary materials. Plasmids encoding LARP-I (pZB578; pZB648) and LARP-I5CHO (pZB580) are available for noncommercial users through AddGene (**<u>accession numbers to be provided upon publication</u>**). E. coli strains are available upon request through a UBMTA.

## Supplementary Materials

Materials and Methods

Figs. S1 to S13

Table S1

References (*54*-71)

