## Supplementary Information for "Dynamic regulation of engineered T7 RNA polymerases by endogenous metabolites"

***This supplementary materials contains:***

Materials and Methods
Figs. S1 to S13
Table S1
References (*54–71)*

#### *Materials and Methods*

##### Reagents

All reagent sources with catalog numbers are provided in the "Reagents" tab of Supporting Data 1.

##### Plasmid and plasmid library generation.

A complete list of plasmids, libraries, gene blocks, and primers are in **Supplementary Data 1**. This list also contains Benchling files for exemplar sequences. All sequences were ordered as synthetic dsDNA gene blocks (eBlocks/gBlocks, IDT) or were PCR amplified using gene-specific primers (IDT) and commercially available kits. Plasmids were constructed by restriction cloning, Golden Gate assembly [1], Gibson assembly [2] or DNA HiFi MasterMix using commercially available kits, by site-directed mutagenesis using commercially available kits (Agilent QuikChange II or NEB Q5®), or by combinatorial nicking mutagenesis [3,4]. Most plasmids were sequence verified using Oxford Nanopore (Plasmidsaurus). All other plasmids had sequence verification of the inserted T7 RNAP using Sanger sequencing (Genewiz).

The base T7 RNAP plasmid pZB017-T7-His6-WT encoding T7 RNAP K378R with an N-terminal 6x-His Tag (MRGSHHHHHHGS-) was constructed from pT7-911Q-His6-WT (Rio 2013), a gift from Seelig Lab at the University of Minnesota. The His tag and K378R mutation appears to have no impact on the activity of the T7 RNAP in benchmark comparisons with commercially purchased T7 RNAP (data not shown). RNAP and LARP constructs containing a *P. putida* ribosome binding site and ATG start codons were restriction cloned into pJH0204, and the T7-sfGFP reporter was cloned into pJH0228, essentially as described [5].

Construction of the T7 RNAP mutational library was previously described [1]. We used the high resolution structures available for T7 RNAP (PDB codes: 1CEZ, 1QLN, 1H38, 1MSW, 3E2E) [6–9] to identify mutations with (i.) any heavy atom within 6 Å of the Cb of position 727; and (ii.) Ca-Cb vector pointing towards position 727. The following sets of mutations were encoded in the library - 725AGV; 727G; 735AGV; 737CDFGHILNQSVY; 778ILV; 781HKNQRS; 782CDFGHILNSVY; 785AGST; 786DEGHKNQRS & 845CDFGHILNSVY.

##### Strain development

Constructs were integrated into engineered genomic sites (serine integrase attP arrays) in *P. putida* AG4775 essentially as described[10] by co-transforming the Bxb1 or the TG1 recombinase with the construct in pJH0204 or pJH0228, respectively. Transformations were performed by electroporation.

The non-essential tryptophanase gene tnaA was knocked out of the *E. coli* B1H US0 strain yielding US0 pyrF-/hisB-/tnaA-, as well as the *E. coli* phage infection strain MG1655 yielding MG1655 tnaA-. To knockout the indole-producing tryptophanase gene from both strains, a Cas9 recombineering protocol was used as described by Choudhury et al and Morgenthaler et al [11,12]. The genomes of *E. coli* BL21, K-12, and W were screened for appropriate gRNA sequences around the *tnaA* locus in the *tnaCAB* operon (Strain, Genomic Location, Accession Code; BL21, 3746691-3748106, CP010816.1; K-12, 3886753-3888168, U00096.2; W, 4111028-41112443, CP002185). Two NGG containing gRNA sequences in the middle of the tnaA gene (ACCATCACCAGTAACTCTGC; ctggctcaataacacgaatg) were selected. Additionally, the DNA used for recombination (gBlock 21) was designed to remove 1032 bp, or more than 70% of the tnaA gene sequence. A genome PCR and Kovak's reagent indole test were conducted to verify successful knockout of the tnaA gene.

Genome PCR was performed by picking a single colony into 1 mL of LB and growing to an $OD_{600}$ ~1. The culture was then pelleted at 17,000xG and washed three times by resuspending in 1 mL nuclease free water and re-centrifuging. After the final wash and resuspension, cells were sonicated for 10 minutes. A 20 uL PCR was performed using primers at 1 uM [407, 408; 407, 409; 407, 410], 6 uL of the sonicated cells, and NEB Q5 MasterMix. The products from the genome PCR were ran on a 2wt% agarose-TAE gel with SYBR Safe, visualized with a SaferImage plate reader, and compared with the products from WT cells.

Kovak's reagent was prepared (2 g p-dimethylamino-benzaldehyde, 10 mL 37% HCl, 30 mL Isoamyl alcohol) and mixed at 50 $\mu$L in 2 mL of a saturated bacterial culture and observed for the formation of pink coloring. No pink color was observed in the selected knockout strains. Successful curing of temperature sensitive plasmids was verified by replica plating on LB-agar with and without chloramphenicol or ampicillin.

**Protein Expression and Purification**

Plasmids containing T7 RNAP proteins driven by a lac-inducible promoter were transformed into chemically competent *E. coli* BL21, grown at 37°C in LB supplemented with 1 (w/v)% dextrose and appropriate antibiotic. At an $OD_{600}$ =[0.6-1.0] the protein expression was induced with 0.25 mM IPTG and allowed to express for 4 hours in a shaker for 6 hours at 30 °C (variants for data reported in **Supp Fig 2**, **Supp Fig 4**), or for 16 hours at 18 °C (all other data). After induction, cultures were immediately spun down at 4000xg, and stored at -80 °C until purification. Proteins were purified by Ni-NTA affinity chromatography essentially as described by Rio et al [13] using BPER and Turbonuclease for lysis. Protein yields ranged from 5-50 mg/L of culture volume. Proteins were stored at 4 °C T7 storage buffer (50mM Tris-HCl pH 7.9, 1mM EDTA, 5mM bME, 100mM NaCl, 15% glycerol) until use.

**Transcriptional Assay**

Transcriptional activity was tested using an adaption of a previously described fluorescent microtiter plate assay [14] . Reactions were assembled in a Black Costar Round-bottom 96-well plate. A reaction mixture (90 µL total) and the T7 RNAP (10 µL total) were mixed and incubated at 37 C and the increase in fluorescence measured. The 100 µL reaction mixture consisted of 20 µL of 5x Transcription Buffer (640 mM HEPES pH 7.5, 112 mM $MgCl_2$, 200 mM DTT, 6.4 mM Spermidine), 20 µL of 25 mM rNTP Mix, 10 µL 1 M KCl, 50 ng pT7-Spinach Template DNA (111 bp) or pT7-8Peppers (555 bp), 1 µL 10 mg/mL BSA, 1 mL 0.1 U/mL iPPase , 2 µL 100 µM 3,5-difluor-4-hydroxybenzylidine imidizolanone (DFHBI) (in ethanol) or 1 µL 100 µM HBC530 (in DMSO) , 2 µL 1 (w/v)% Triton X-100, and a balance of Nuclease Free Water to 90 µL. This assembled reaction mixture sans enzyme was added to a Black Costar Plate Well containing 10 µL of 20 µM Protein and mixed by pipetting. The fluorescence readout (Spinach: Ex/Em 469/501; Peppers Ex/Em 485/530) was performed every 60-72 sec for at least 60 minutes in a Biotek Synergy H1 plate reader set to 37 °C. Transcriptional activity was determined as the slope of the increase in fluorescence (RFU/min). RFU/min was calculated by fitting a line to a moving window of 20 data points, and reporting the largest value. For low activity, a correction to the max rate is used where the slope is multiplied by the correlation coefficient of the fit, and the highest value is reported.

***In vivo* selection for identification of LARPs**

The T7 RNAP screen was developed from a previous selection used for screening Zinc finger nuclease target sequences [15,16]. An auxotrophic strain, US0 pyrF-/hisB-, containing the homologous HIS3 and URA3 deletions was complemented with a plasmid containing the HIS3/URA3 genes under a pT7 promoter. Additionally, the gene for T7 RNAP was transformed under a secondary plasmid. Initial tests of growth screening on solid media suggested the pT7/T7 RNAP system was toxic, as previously documented [17,18]. To reduce the toxicity of T7 RNAP, different pT7 promoter strengths and T7 RNAP expression strengths were tested. Two plasmids were co-transformed into US0 pyrF-/hisB- and plated on M9-Complete (M9 mineral salts with 6 g/L dextrose, 1.4 g/L Mix Amino Acids (-Trp, -Leu, -His, -Ura), 78 mg/L Trp, 22.4 mg Uracil, 380 mg Leucine, 0.5 g/L of Histidine and 0.2 g/L Yeast Extract) 1.8 (w/v)% agar plates with Tetracycline (10 $\mu$g/mL), Chloramphenicol (30 $\mu$g/mL), and Kanamycin (50 $\mu$g/mL) and grown overnight at 37 °C. Fresh colonies were selected the next day into M9-his (M9 mineral salts with 6 g/L dextrose, 1.4 g/L Mix Amino Acids (-Trp, -Leu, -His, -Ura), 78 mg/L Trp, 22.4 mg Uracil, 380 mg Leucine) liquid culture and grown overnight. Overnight cultures were diluted to an $OD_{600}$ of 0.01-0.04 into 3.5 mL of M9-his with 1 mM of the HIS3 competitive inhibitor 3-aminotriazole (3-AT). Growth was measured by $OD_{600}$ in Hungate tubes as described [19]. The ultimate screen used in this work had a lacUV5 promoter driving expression of full length T7 RNAP with a GUG initiation codon and the UmuD N-degron tag [18,20].

For library selections, 1800 ng of the previously developed library [1] was transformed into prepared electrocompetent cells of the selection strain (US0 pyrF-/hisB-/tnaA-) already containing the pT7-H3U3 plasmid (pZB260), resulting in 20E7 estimated transformants (range 10E7-31E7). The transformation was directly plated on large Corning bioassay (245x245mm) plates with M9-Complete media. The resulting libraries were scraped with M9-Complete liquid media and prepared as glycerol stock (15 (w/v)% glycerol). The negative selection was performed by plating approximately 5E9 cells (10 OD-mL) on agar containing M9-Complete media and 1 mM 5-fluoroorotic acid (5-FOA), and incubated at 37°C overnight. After negative selection plates were scraped with M9-his, washed two times by pelleting (4000xG for 10 mins) and resuspending in M9-his, and either directly used for positive selection or prepared as glycerol stocks and stored at -80 °C.

For positive selection, M9-his plates were prepared with 1 mM 3-amino-1,2,4-triazole and either 50 $\mu$M of indole or indole-5-carboxyaldehyde or ethanol as the solvent control. Cells passed through counter selection were plated on the selective media plates and grown for 20 hours at 37°C. The resulting colonies for each selection condition were scraped with M9-his media and prepared as glycerol stocks. DNA from resulting libraries was plasmid extracted and prepared for next generation sequencing essentially following method B of Kowalsky et al [21]. The libraries were sequenced on an Illumina MiSeq using 2 × 250 paired end reads by Rush University genomics core.

**Next-generation sequencing**

Raw reads from the sequencing were analyzed essentially as described in Smith et al [22]. Briefly, fastq files were merged using FLASh [23]with standard inputs. Merged sequences were then converted to fasta files. Next, sequences were sorted by correct length (416 bp) and absence of any ambiguous base-calls (e.g. "N" in sequence). Sequences that passed filtering were translated to an amino acid string using Biopython [23]and screened for a proper initial sequence of three amino acids. Last, an identifying mutational string was generated for each read constrained to only encoded library positions and the prevalence of each mutational string was counted and divided by all observed sequences to obtain frequencies. These frequencies were then used in the subsequent PSERM analyses.

**Planktonic Clonal Growth Assays**

For clonal growth assays, LARP plasmids were freshly transformed into the selection strain containing the pT7-H3U3 plasmid (US0 pyrF-/hisB-/tnaA-; pZB260) and dilution plated on M9-C (+ 1 mM 5-FOA). Individual colonies were selected from M9-Complete (+ 1 mM 5-FOA) and grown overnight in M9-his media. The following morning, cultures were diluted 50 $\mu$L into 3.5 mL of M9-his (+3-AT) containing 50 $\mu$M ligand or ethanol as the solvent control in 14 mm Hungate tubes resulting in starting $OD_{600}$ of 0.01-0.04. Hungate tubes were incubated aerobically at 37 $^{o}$C and shaking, and the $OD_{600}$ was measured every 30-90 minutes for up to 8 hours or until cultures reached saturation ($OD_{600}$ = 0.6).

**GFP Reporter Assays**

For exogenous activation experiments, colonies were incubated in 400 $\mu$L of SOB containing the appropriate antibiotic and grown overnight for ~16 hours. Saturated cultures were then diluted 40 $\mu$L into 360 $\mu$L of fresh SOB containing the appropriate antibiotics and indole concentrations (or ethanol as a vehicle control) and grown for an additional 22 hours prior to measuring GFP. For endogenous activation experiments, colonies of each variant were picked into 400 $\mu$L of SOB containing the appropriate antibiotic and grown overnight. Saturated cultures were then diluted 40 $\mu$L into 360 $\mu$L of fresh SOB containing the appropriate antibiotics and indole concentrations (or solvent control) and grown for an additional 22 hours prior to measuring GFP. For both experiments, GFP (ex: 485 nm; em: 515 nm) was measured with 200 uL of undiluted cultures in 96 well black costar round bottom plates (REF# 3792) and the pathlength corrected OD600 was measured at 200 $\mu$L total volume in 96 well clear flat bottom plates (fisherbrand; Cat. No. 12565501) following a 5x dilution into SOB. All measurements were performed using a Biotek Synergy H1 plate reader at room temperature and automatic gain.

For co culturing experiments, colonies of each variant (all senders expressing mRFP; all receivers containing pT7-sfGFP as pZB604; sender: US0 $tnaA^+$; sender control: US0 $tna^-$; LARP-I expressing receiver: US0 $tnaA^-$; T7 $RNAP^{R632S}$ expressing receiver positive control: US0 $tna^{+/-}$; T7 $RNAP^{R632S/Y639A}$ expressing receiver negative control: US0 $tna^{+/-}$) were picked into 400 $\mu$L of SOB containing the appropriate antibiotic and grown overnight for ~16 hours. Saturated cultures were then measured for $OD_{600}$, spun down, resuspended in fresh SOB containing the appropriate antibiotics, and mixed at 1:1 ratio of OD-mL to obtain approximately equivalent cell populations. Samples were then grown for 22 hours prior to measuring RFP/GFP fluorescence and the 5x dilution of the $OD_{600}$ at 200 $\mu$L volume. The non-diluted samples were used for imaging (RFP Ex: 583 nm; Em: 609 nm).

**Fluorescence imaging**

Fluorescent images of the co-culture experiments reported in **Fig 2** used 5 $\mu$L of diluted culture ($OD_{600}$ = 0.1) plated on a 2 cm x 1 cm section of minimal media plates (M9) with 1.5 (w/v) % agarose. After cells dried, the agarose was inverted onto an ibidi $\mu$Slide well plate (Cat. No. 80427) and imaged on an Olympus Microscope. Brightfield was used to find cells that were of suitable imaging density. Exposure time and fluorescent channel were manually set, and then the laser was opened (20% laser power). Exposure time was 2 ms for Brightfield, 100 ms for GFP channel (Ex: 488 nm; Em: 520 nm), and 300 ms for RFP (Ex: 594 nm; Em: 610 nm) channel. Images were taken in the first few seconds following laser exposure and magnification adjustment to reduce bleaching. Images were processed in ImageJ [24] by combining the individual channels (BF, GFP, RFP), adjusting brightness and contrast for the entire channels to reduce over-exposure, (BF-gray minimum/maximum 2700/4000, RFP-red/GFP-cyan minimum/maximum 100/4000). Representative images are shown in Figure 2F of the paper, while all data can be accessed in **Source Data**.

**Phage Assays**

Phage were amplified in 50 mL cultures of LB and titered following Kibby et al. [25] and stored at 4$^{o}$C following filtration through a 0.22 $\mu$m filter. For gp1 deficient phage strains, *E. coli* BL21*(DE3) was used to amplify the phage. Phage was diluted in the same LB media used for amplification.

To perform phage plaque assays, MG1655 tnaA- was transformed with pZB513 (plasmid expressing WT T7 RNAP) and pZB578 (LARP-I). Single colonies were picked and grown overnight at 37 $^{o}$C in LB + 30 $\mu$g/mL chloramphenicol. The next day, LB overlay plates were prepared following Kibby et al [25]. Briefly, 3 mL of melted 1.5 (w/v)% LB-agar was added to 6 mL LB containing 30 $\mu$g/mL chloramphenicol, phage infection salts (10 mM $MgCl_2$, 10 mM $CaCl_2$ and 100 $\mu$M $MnCl_2$), and indole or ethanol as a vehicle control. To this mixture 100 $\mu$L of saturated overnight culture was added, immediately mixed, and plated on top of pre-prepared LB plates. The overlay agar was allowed to dry for at least 15 minutes and not more than 1 hour at room temperature. After plates were dried, 4 $\mu$L of phage serial dilutions were plated onto the overlay and allowed to dry. Plates were then covered and placed in the static incubator at 30 $^{o}$C for 14 hours.

To perform phage clearance in planktonic cultures, single colonies of MG1655 tnaA- containing pZB513 (WT T7 RNAP) or pZB578 (LARP-I) were picked and grown for 6-8 hours in LB containing chloramphenicol and phage infection salts. Cultures were back-diluted to an $OD_{600}$ of approx. 0.2 in LB containing 30 $\mu$g/mL chloramphenicol, phage infection salts (10 mM $MgCl_2$, 10 mM $CaCl_2$ and 100 μM $MnCl_2$), and indole or ethanol as a vehicle control. Phage infection was carried out in 96-well plates with 200 $\mu$L total volume per well by adding 4 $\mu$L of phage dilutions at a multiplicity of infection (PFU/cell, assuming 5E8 *E.coli* per OD-mL) of 0.5 or 5.

##### *P. putida* growth experiments

*P. putida* colonies with stable integrations of the T7 reporter and T7 RNAP or LARP constructs were grown aerobically overnight at 30°C, shaking at 250rpm, in LB with kanamycin (50 µg/mL) and streptomycin (50 µg/mL). Cultures were back-diluted to an $OD_{600}$=0.05 in fresh media, then indole (or ethanol as a vehicle control) was added at the concentrations indicated. Cultures were grown at 30°C, shaking at 250rpm prior to measurement of $OD_{600}$ and GFP by flow cytometry. Induction of sfGFP was measured at 24 h of growth in indoles by flow cytometry. Per-cell GFP were quantified using the sorting gates shown in **Supplementary Figure 11**.

##### Promoter Specificity Testing

Orthogonal polymerase-promoter specificities were tested by cloning the different promoter sequences in frame of the canonical pT7 sequence in pZB260, upstream of the HIS3/URA3 genes resulting in pH3U3 promoter plasmids (pZB612: pCGG; pZB613 pCTGA,pZB616: pN4). Additionally, corresponding specificity loop mutations for T7 RNAP were incorporated into the pB1H1-LARP-I plasmid (pZB578), resulting in plasmids (pZB643-LARP-I-pCGG, pZB644-LARP-I-pCTGA, pZB647-LARP-I-pN4). All promoter and polymerase combinations (16) were transformed into the selection strain and plated on non-selective media. Fresh colonies of each combination were picked into 400 $\mu$L of M9-Complete media and grown for 12 hours to saturation ($OD_{600}$ ~1.0). Cultures were diluted 100x into M9-his (+ 1 mM 3AT) and 5 $\mu$L was plated on solid selective media M9-his (+1 mM 3-AT; +/- 100 $\mu$M indole) and counter-selective M9-Complete media (+1 mM 5-FOA; +/- 100 $\mu$M indole). Plates were incubated at 37$^o$C for 18 hours before imaging.

##### Circular dichroism

Circular dichroism was performed on an Applied Photophysics Chirascan V100. Purified LARP-I was buffer exchanged into 20 mM NaPO4, pH 7.8 with 50 mM NaCl using 7 kDa Zeba 0.5 mL spin columns per manufacturer's protocol and diluted to a concentration in the range of 0.1-0.4 mg/mL. 170 $\mu$L of sample was loaded into clean 0.5 mm cuvettes. Due to the presence of 50 mM NaCl, the usable spectra were limited to 195 nm. Spectral measurements were taken using 1 sec timepoints at 2 nm bandwidth and 0.5 nm step size for LARP-I in the presence and absence of 500 $\mu$M indole. The ellipticity spectra of LARP-I from 195-260 nm was indistinguishable from TS at both 0 and 500 uM indole concentrations. For CD melt curves, the wavelength was maintained at either 208.5 nm or 222 nm and the ellipticity was measured for 24 secs between 0.5 C temperature steps as the sample was heated at 1 $^o$C/min.

##### Fluorescence Anisotropy

The canonical 17-nt T7 promoter (5'-TAATACGACTCACTATA-3') spanning from -17 to -1 position of the initiation start site was purchased from IDT with 5' 6FAM-fluorescein conjugated to the 5' end of the forward promoter sequence [26]. The fluorescent promoter was annealed at 20 $\mu$M concentration in buffer containing 10 mM Tris, pH 7.8, 50 mM NaCl, and 1 mM EDTA with the complementary strand in 20% excess (24 $\mu$M) by heating the mixture to 98 °C for 1 minute and cooling to 4 °C over 3 hours in a thermal cycler. Annealed primer was diluted to 2 nM in a buffer containing 33 mM HEPEs, pH 7.8, 50 mM NaCl, and 10 mM MgCl2. Protein was exchanged into 33 mM HEPES, pH 7.8, 50 mM NaCl, and 10 mM MgCl2 using 7 kDa Zeba 0.5 mL spin columns per manufacturer's protocol. Protein concentration was determined using A280 nanodrop as well as Bradford assay. Protein was serial diluted 22 times in 2x dilutions (1:1) or 1.5x dilutions (2:1) in the same buffer. Equivolume mixtures of DNA solution and protein dilutions (DNA final concentration: 1 nM, final volume 20 $\mu$L per well) were prepared in 384 well plates in triplicate. Samples were allowed to incubate at 25 °C or 37 °C for at least 1 hour prior to measurement. Fluorescence polarization was measured using a Tecan Spark plate reader with automated settings for FAM-fluorescein channels (excitation wavelength of 483 nm; excitation bandwidth of 20 nm; emission wavelength of 529 nm; emission bandwidth of 20 nm; automatic gain; mirror: automatic -dichroic 510; 30 flashes; integration time: 40 $\mu$s; settling time: 100 ms; z-position: 20000 $\mu$m; G-factor calibrated from a buffer blank and reference sample of 1 nM fluorescent DNA in buffer) [27].

Analysis of the fluorescence anisotropy data was performed using a Python package (https://github.com/nicklammer/AnisotropyBindingFit) to fit data to a 1:1 binding isotherm[28].

### FIGURES

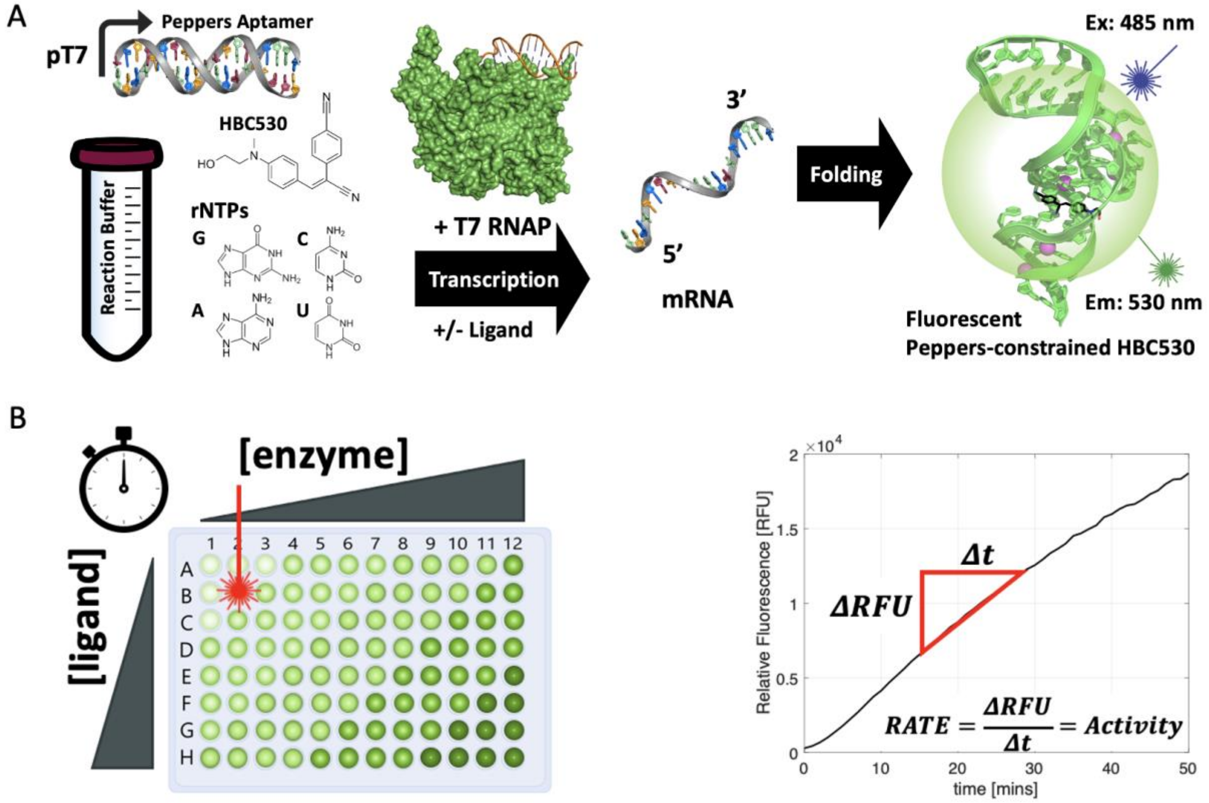


***Supplementary Figure 1. Overview of in vitro transcriptional activity assay***. ***[A]*** *A transcriptional assay is assembled using a dsDNA sequence encoding an aptamer (i.e. Peppers or Spinach) under the pT7 canonical promoter sequence 5’-TAATACGACTCACTATA-3’) in solution with reaction buffer, rNTPs, the non-fluorescent aptamer ligand (i.e. HBC530 or DFHBI). The reaction is initiated with the addition of T7 RNA Polymerase, at various ligand concentrations. The RNA polymerase transcribes the mRNA aptamer sequence, which binds to and constrains a chromophore, resulting in fluorescence.* ***[B]*** *The assay is performed in 100 $\mu$L reaction volume in 96 well plates and the output are read via a plate reader with the corresponding excitation wavelength and emissions spectra (HBC530: 485 nm/530 nm; Spinach 24-2: 469 nm/501 nm). Fluorescence over time is used to define activity [RFU/min].*

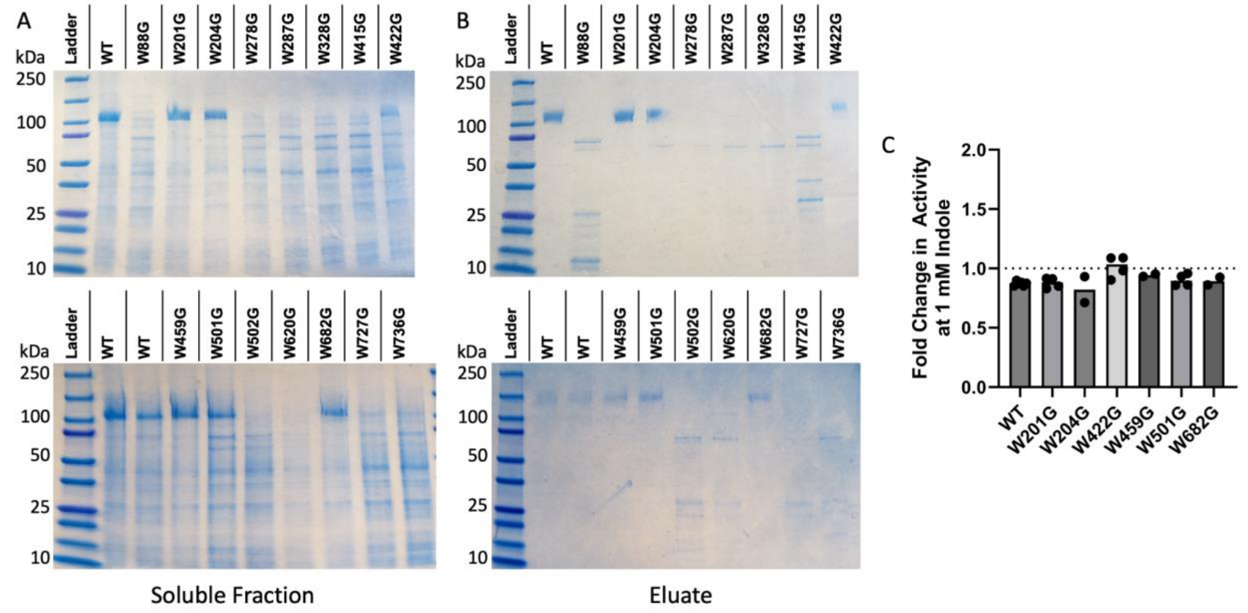


***Supplementary Figure 2. Initial glycine scan of 15 rationally selected tryptophan residues, expression and purification results. [A]*** *Soluble Fraction and* ***[B]*** *Eluate from mutants of T7 RNAP harboring the given mutations. WT: N-terminal $His_6$ T7 RNAP R378K.* [**C**] *Fold-change in activity of initially purified variants in the presence of 2 mM indole relative to the solvent control (ethanol) revealed no variants with positive indole modulation.*

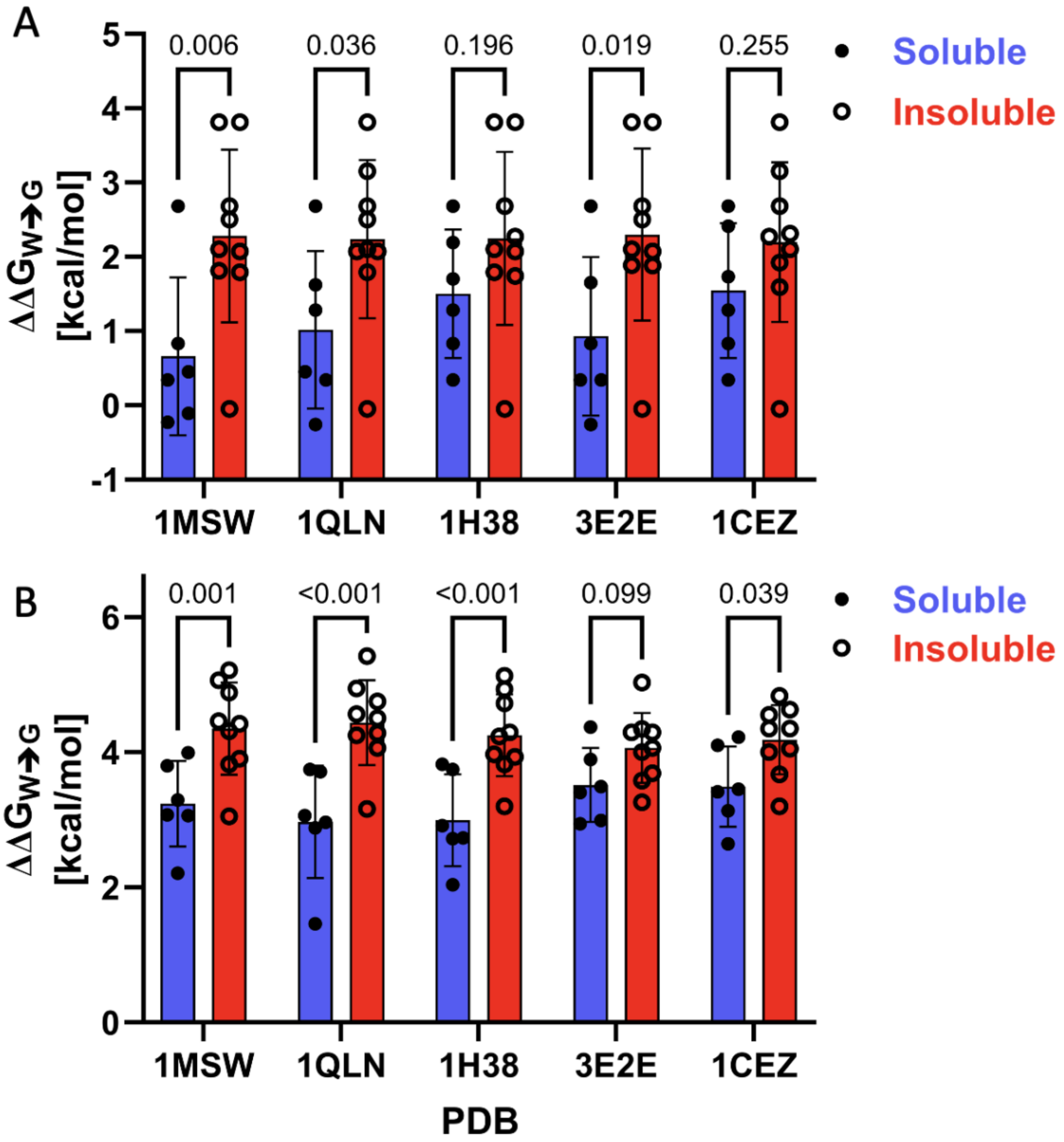


***Supplementary Figure 3. Computationally predicted stability of glycine scan mutants.*** *Two web-based servers* ***[A]*** *SDM2 and* ***[B]*** *PoPMuSiC2.0 were used to ascertain the stability of tryptophan to glycine mutants in the soluble (n=6) and insoluble (n=9) groups from expression results. Five PDB structures were used as inputs, and the p-value for each PDB comparison is listed. For both SDM2 and PoPMuSiC2.0, the insoluble group had a higher destabilizing value than did the soluble group, which was statistically significant ($p<0.05$) for seven of the ten tests. P values were computed by multiple unpaired t tests assuming a single pooled variance and no correction for multiple comparisons.*

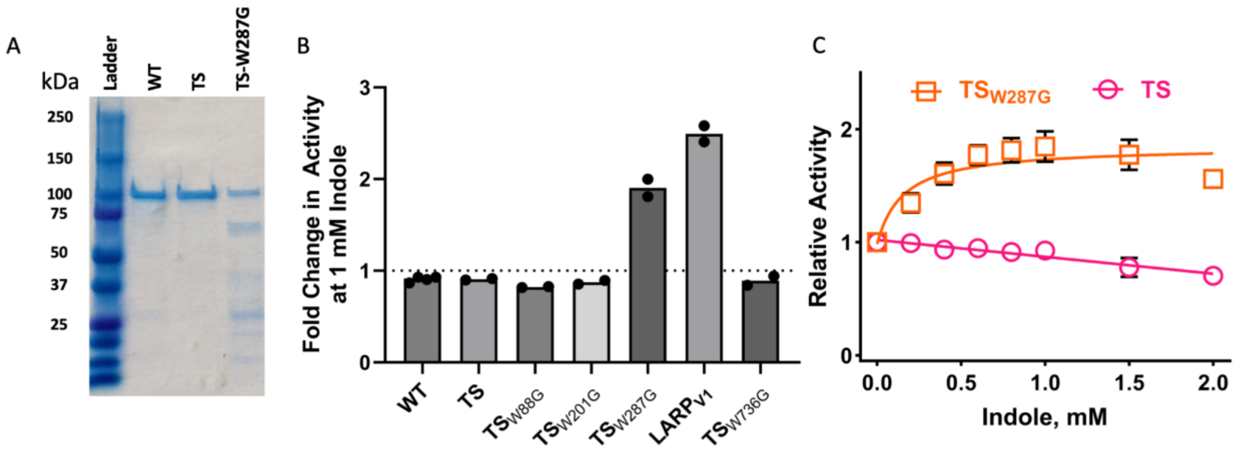


***Supplementary Figure 4. Identification of initial LARPs using a thermally stable in vitro glycine scan.*** *A. Expression of thermally stabilized T7 RNAP TS (S430P, N433T, S633P, F849I, F880Y) and TSW287G. B. Fold-change in activity of tryptophan to glycine mutants in the thermally stabilized background in the presence of 2 mM indole relative to the solvent control (ethanol) revealed two variants - W287G & W727G ($LARP_{V1}$) - which had ~>2x increase in activity at 2 mM indole. Individual data points indicate biological replicates. C. Dose-response curve for TSW287G and TS. Activity is reported normalized to 1 for each variant in the absence of indole.*

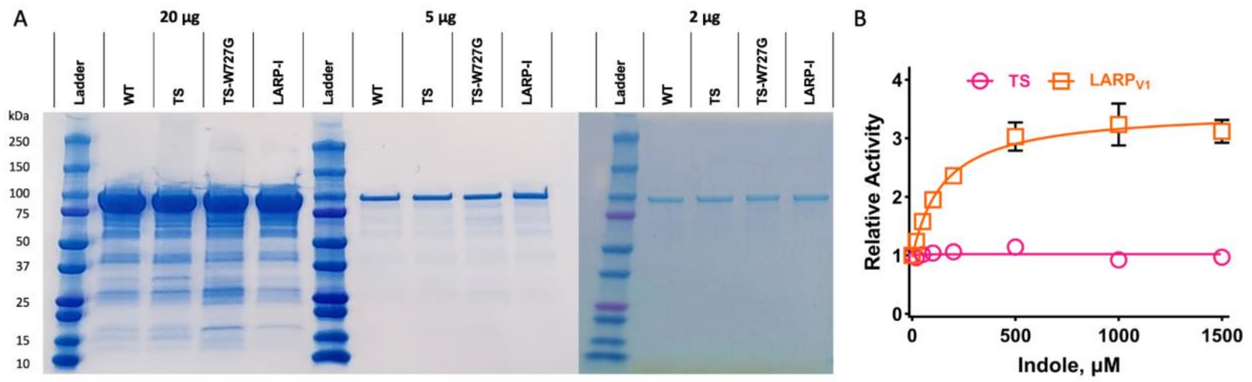


***Supplementary Figure 5. Activity of LARPv1 (TS-W727G). A.*** *SDS-PAGE Protein Gel of purified wild-type (WT), thermally stable (TS), TS-W727G (LARPv1), and LARP-I at 20 µg, 5 µg, and 2 µg.* ***B.*** *Dose-response curve for TSW287G and TS. Activity is reported normalized to 1 for each variant in the absence of indole.*

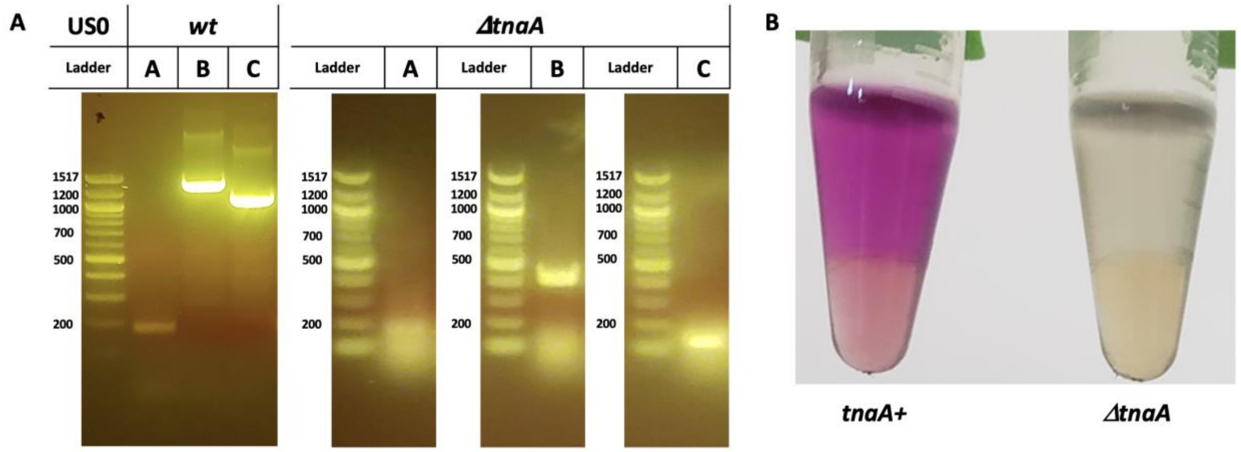


***Supplementary Figure 6. Generation of E. coli US0 pyrF-/hisB- ΔtnaA by CRISPR-Cas9.*** *A CRISPR-Cas9 dsDNA break and homology directed recombination protocol utilizing temperature-sensitive plasmids was used to generate a knock-out of the gene encoding the tryptophan degrading enzyme tryptophanase (tnaA). A gRNA targeting the tnaA coding sequence and a linear fragment of dsDNA with homology arms surrounding the tnaA coding sequence were used to remove a large section of the E. coli genome in the selection strain US0 pyrF-/hisB-.* ***[A]*** *Successful knockout of the tnaA gene was confirmed through colony PCR of the tnaA gene. Three pairs of primers amplifying the genomic region of tnaA subject to the ~1032 bp genomic KO knockout were tested. Region A: 191 bp in WT, N/A in tnaA KO; Region B: 1452 bp in WT, 420 bp in tnaA KO; Region C: 1156 bp in WT, 136 bp in tnaA KO.* ***[B]*** *A phenotypic test using Kovac's reagent on saturated cultures confirmed indole is not produced in the ΔtnaA strain. Kovac's reagent reacts with indole, producing a brilliant purple color.*

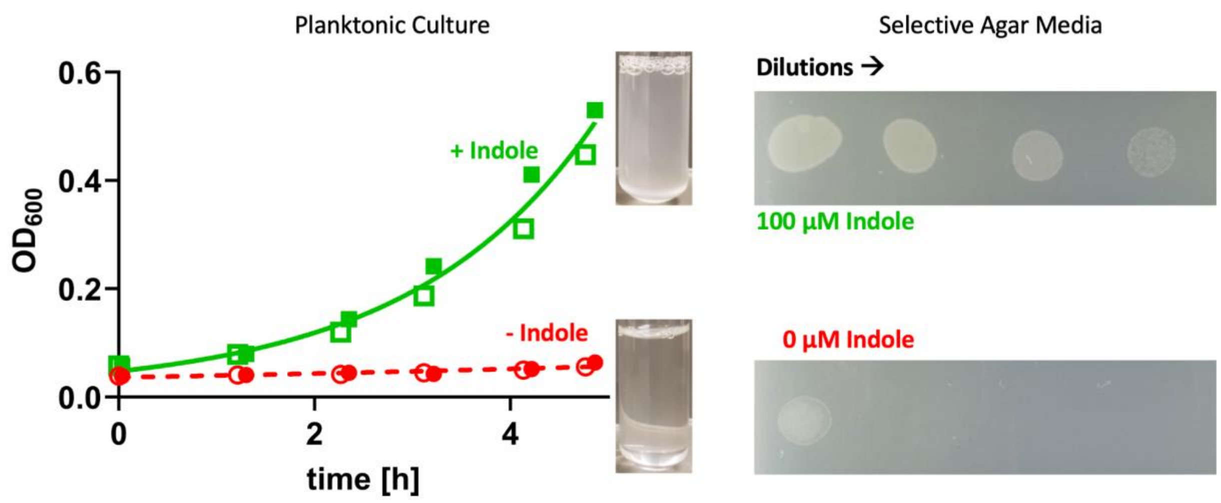


***Supplementary Figure 7.*** *Optimized T7 positive and negative selection assay affords indole-dependent growth for LARPv1. The left panel represents growth in planktonic culture of E. coli US0 pyrF-/hisB- ΔtnaA expressing LARPv1 and HIS3/URA3 driven by a strong T7 promoter. Open and closed symbols represent different biological replicates. + indole is a condition with 100 µM indole. The right panel represents growth on solid agar selective media with and without the indicated concentration of indole.*

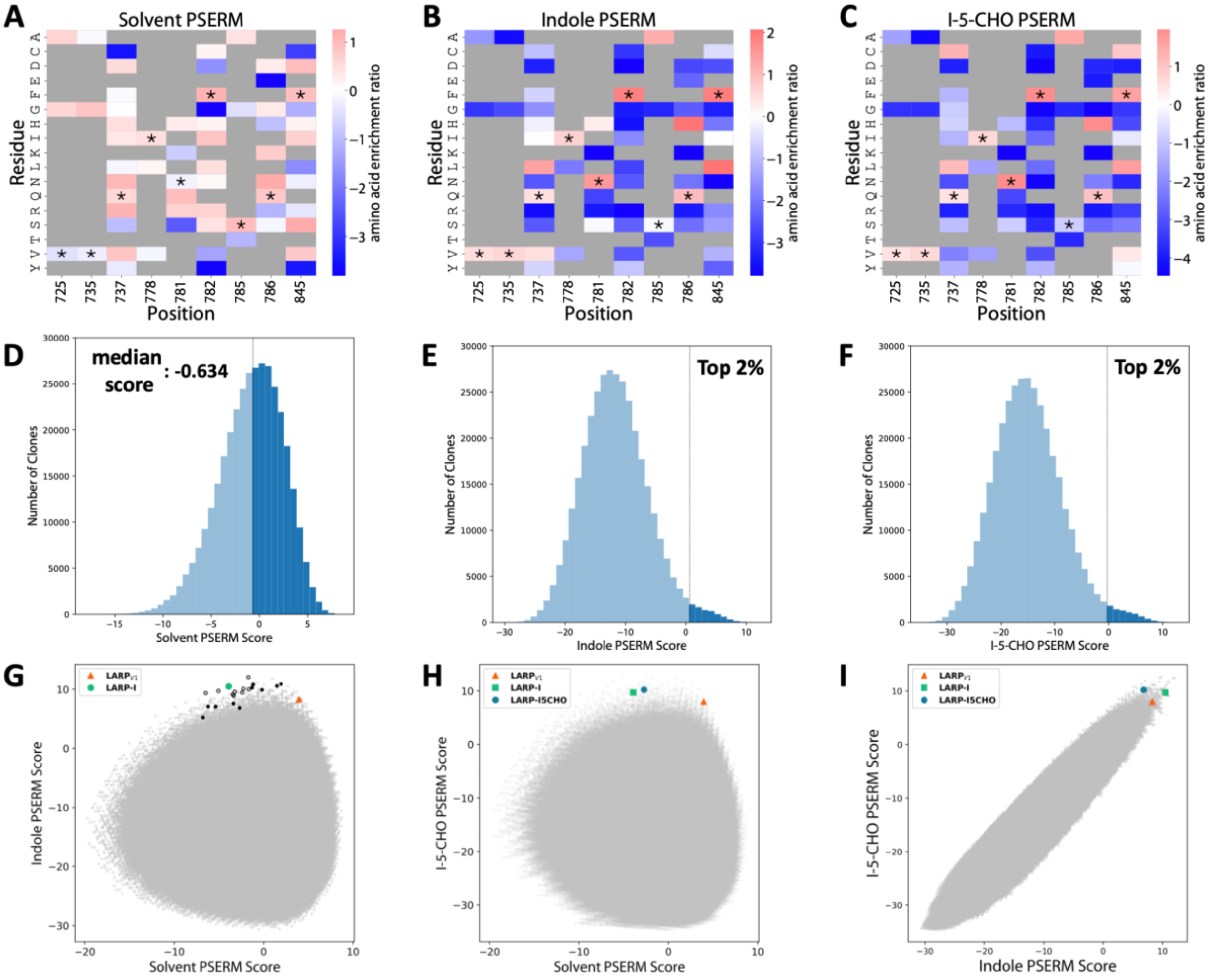


***Supplementary Figure 8. PSERM Scoring [A-C] Position Specific Amino Acid Enrichment Ratio Score Heat Map, [D-F] PSERM score distribution for library variants, [G-I] relevant PSERM scores plotted against each otherfor [A] Solvent control, [B] Indole, and [C] Indole-5-carboxyaldehyde Selection Results.*** *PSERM scores for each variant result from the additive score of constituent mutations pictured for Solvent [A], Indole [B] and Indole-5-carboyxaldehyde [C]. WT residues are flagged with an asterisk (*).The top 2% of library variants in the ligand selections (E, F) have a positive PSERM score. Plotting the scores against each other reveals Pareto-fronts in the Solvent vs Ligand plots (G, H) and high correlation between Indole and Indole-5-carboxyaldehyde scores (I). Larger closed symbols represent hits tested and variants described in the main text.*

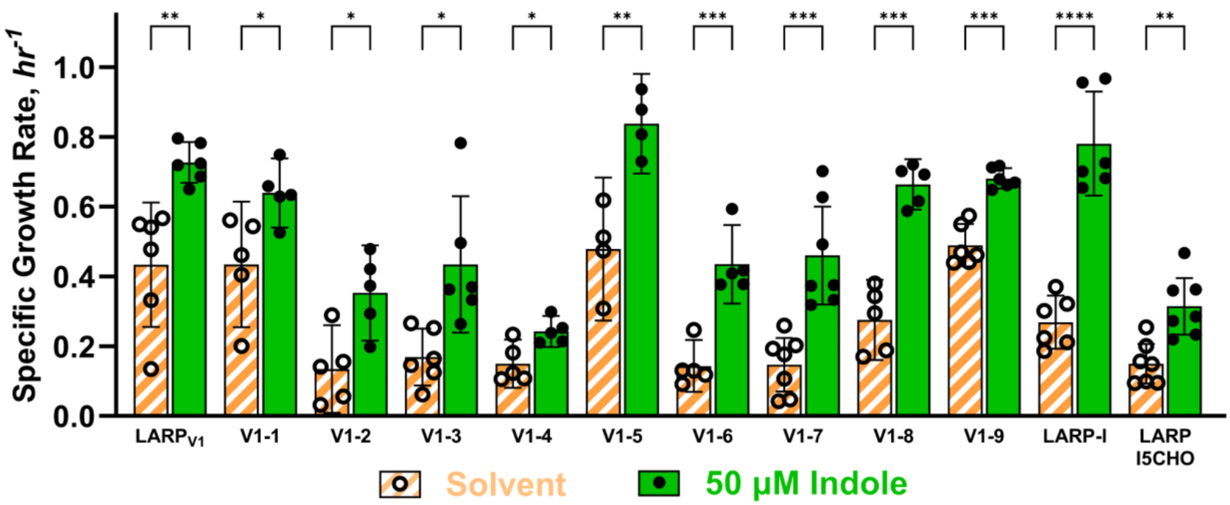


***Supplementary Figure 9.*** *Individual Growth Rates of LARP hits in selective media (histidine-deficient, 1mM 3-AT) measured in planktonic growth assays in presence and absence of indole. p-values: ****: p-value<0.0001; ***: p-value<0.001; **: p-value<0.01; *: p-value<0.05. Amino acid sequences of tested LARPs are shown in Table S1.*

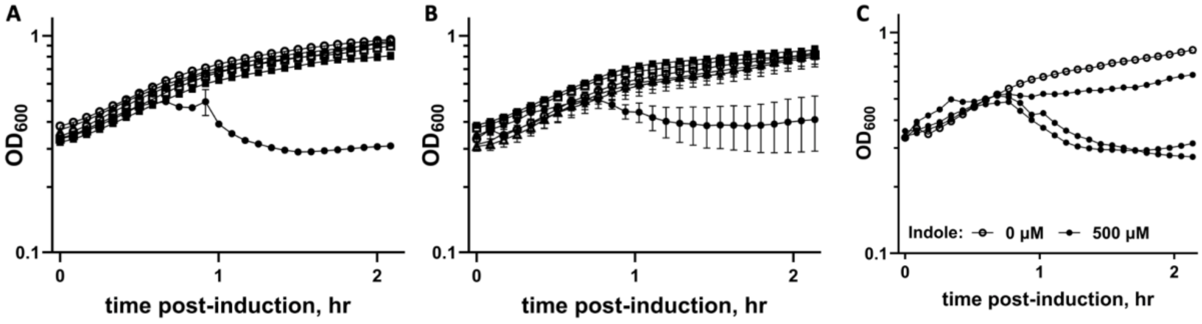


***Supplementary Figure 10. Indole-dependent Sgl$^{KU1}$ expression and lysis in LARP-I containing E. coli.*** *[A] Replicate 1 lysis curves (OD$_{600}$ vs time post-induction in hours) mean and standard error of the mean of biological replicates (n=3) for negative controls (LARP-I, no Sgl$^{KU1}$; catalytically dead T7 RNAP$^{R632S;Y639A}$, Sgl$^{KU1}$) and LARP-I, Sgl$^{KU1}$ with 0 uM indole (empty symbols) or 500 μM indole (filled symbols). [B] Replicate 2 lysis curves (OD$_{600}$ vs time post-induction in hours) mean and standard error of the mean of biological replicates (n=3) for negative controls (LARP-I, no Sgl$^{KU1}$; catalytically dead T7 RNAP$^{R632S;Y639A}$, Sgl$^{KU1}$) and LARP-I, Sgl$^{KU1}$. [C] Replicate 2 experimental group (LARP-I + Sgl$^{KU1}$) with 0 μM indole (empty symbols, Mean and SEM, n=3) or 500 μM indole (filled symbols, individual traces).*

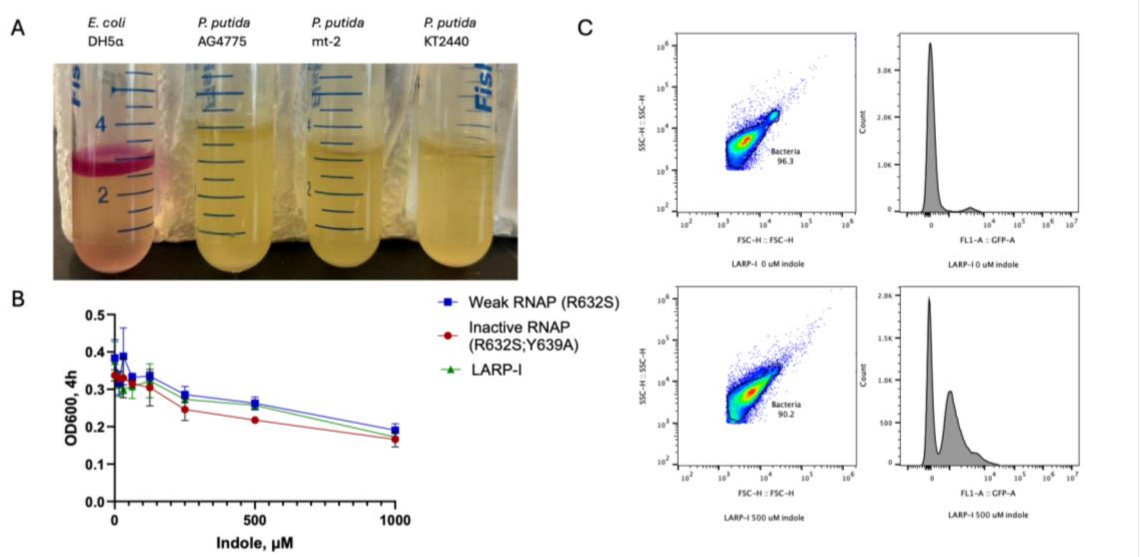


**Supplementary Figure 11.** *Development of LARP-I expressing P. putida strains*. A. Overnight cultures of *E. coli* DH5ɑ, *P. putida* mt-2, and the mt-2 derivatives KT2440 and AG4775 were grown overnight in LB medium before addition of Kovac's reagent. The cherry red color of the Kovac's reagent layer indicates the presence of indoles, as seen for the positive control strain *E. coli* DH5ɑ. B. OD$_{600}$ vs. indole concentration for three *P. putida* strains expressing different T7 RNAP constructs. Cultures were grown aerobically in LB overnight, then back-diluted to an OD$_{600}$=0.05 into indole-containing media at the concentration indicated. OD$_{600}$ was measured 4 hours after indole addition. Higher (500, 1000 μM) indole concentrations leads to impaired growth. Error bars represent 1 s.e.m. (n=3). C. Flow cytometry gating strategy and sample GFP expression histograms in the absence and presence of indole. Data shown is the *P. putida* LARP-I expression strain at 0 and 500 μM indole.

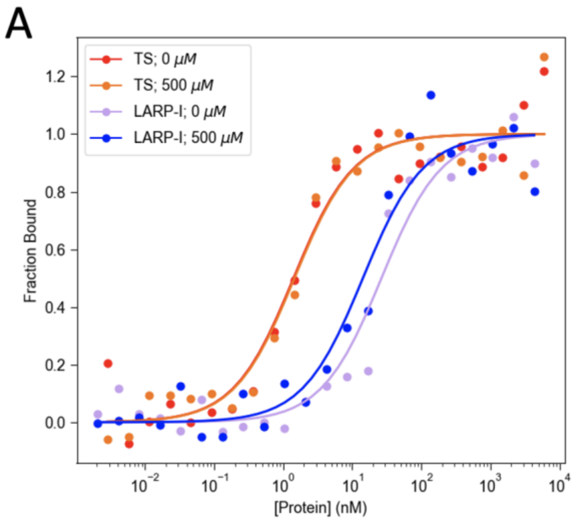


**Supplementary Figure 12.** *Fluorescence anisotropy of purified TS and LARP-I at 1 nM DNA (pT7: -17:-1) at 25 °C [A]and 37 °C [B]. Values are the means of three technical replicates fit to a 1:1 binding isotherm.*

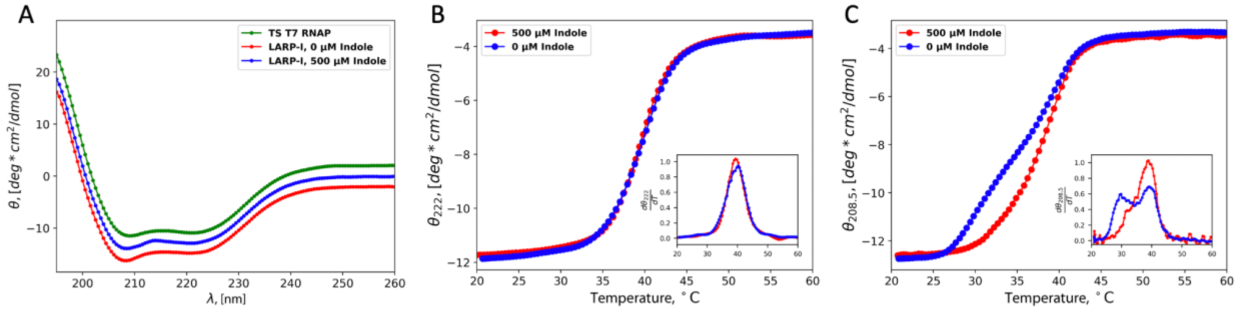


**Supplementary** ***Figure 13. Circular Dichroism Spectroscopy of LARP-I.*** *Full Spectra of LARP-I with and without 500 µM shows no difference relative to the TS background. Circular dichroism temperature ramp of purified LARP-I in the presence or absence of 500 µM indole measuring ellipticity at 222 nm [B]and 208.5 nm [C]. LARP-I shows a decreased* $T_{m,app}$ *around 39°C as measured by the maximum of the derivative of the change in ellipticity (inset). [C] At 208.5 nm LARP-I shows two distinct peaks in the derivative of the ellipticity in the absence of indole (29°C, 39°C), which is less pronounced in the presence of 500 µM indole, suggesting differential stabilization of other secondary structure elements.*

### TABLES

*Table S1. Sequences and properties of tested LARP constructs. Growth rate data indicates that the constructs are inducible under plate conditions and were tested in planktonic culture shown in Fig S9. Blank means whether the variant grew on selective solid media in the absence of ligand (in all cases +: yes, - : no). Negative selection indicates whether the variant grew on counterselective solid media in the presence of 1 mM 5-FOA. Positive selection indicates whether the variant grew under positive selection conditions on selective media in the presence of indoles. The tested variants all had Positive: +; Negative: +; Blank: - (gray shading).*

| | | | PSERM Score | | | Growth | Selection Conditions | | |
|---|---|---|---|---|---|---|---|---|---|
| Name | Mutations | PSERM Designation | Blank | Indole | Lz2 | Rate | Positive | Negative | Blank |
| LARP-V1 | (V725, W727G, V735, Q737, I778, N781, F782, S785, Q786, F845) | VVQINFSQF | 3.947 | 8.265 | 7.969 | + | | | |
| LARP-I | (Q737L, N781S, S785A, Q786H, F845L) | VVLISFAHL | -3.923 | 10.513 | 9.707 | + | | | |
| LARP-I-5-CHO | (Q737C, I778L, S785A, Q786H) | VVCLNFAHF | -2.720 | 6.854 | 10.238 | + | | | |
| V1-1 | (Q737V, S785A, Q786H) | VVVINFAHF | 1.954 | 10.894 | 9.066 | + | | | |
| V1-2 | (Q737L, S785A, Q786H, F845Y) | VVLINFAHY | -3.431 | 7.575 | 10.649 | + | | | |
| V1-3 | (Q737C, I778L, S785A, Q786H, F845L) | VVCLNFAHL | -5.349 | 7.119 | 10.315 | + | | | |
| V1-4 | (Q737C, S785A, Q786H, F845Y) | VVCINFAHY | -6.817 | 5.238 | 10.767 | + | | | |
| V1-5 | (Q737L, N781S, S785A, Q786H) | VVLISFAHF | -1.294 | 10.248 | 9.630 | + | | | |
| V1-6 | (Q737L, N781H, S785A, Q786H, F845L) | VVLIHFAHL | -1.162 | 10.841 | 9.384 | + | | | |
| V1-7 | (Q737C, S785A, Q786H, F845C) | VVCINFAHC | -6.221 | 7.085 | 11.863 | + | | | |
| V1-8 | (Q737L, N781H, S785A, Q786H) | VVLIHFAHF | 1.467 | 10.576 | 9.307 | + | | | |
| V1-9 | (Q737V, N781H, S785A, Q786H, F845L) | VVVIHFAHL | -0.158 | 9.928 | 6.057 | + | | | |
| V1-10 | (Q737L, S785A, Q786H, F845L) | VVLINFAHL | -1.679 | 12.072 | 12.470 | - | + | - | + |
| V1-11 | (Q737C, S785A, Q786H) | VVCINFAHF | -2.436 | 9.470 | 12.510 | - | + | - | + |
| V1-12 | (Q737F, S785A, Q786H, F845L) | VVFINFAHL | -1.643 | 10.004 | 10.020 | - | + | - | + |
| V1-13 | (Q737I, N781S, S785A, Q786H, F845L) | VVIISFAHL | -3.484 | 9.137 | 7.237 | - | - | + | - |
| V1-14 | (Q737H, N781S, S785A, Q786H, F845L) | VVHISFAHL | -3.361 | 8.921 | 7.757 | - | - | + | - |
| V1-15 | (N781S, S785A, Q786H, F845L) | VVQISFAHL | -3.231 | 9.593 | 9.019 | - | - | + | - |
| V1-16 | (Q737L, F782Y, S785A, Q786H, F845L) | VVLINYAHL | -6.494 | 9.367 | 9.184 | - | - | + | - |
| V1-17 | (Q737L, I778V, S785A, Q786H, F845L) | VVLVNFAHL | -2.273 | 10.084 | 10.146 | - | + | - | + |
| V1-18 | (Q737C, S785A, Q786H, F845L) | VVCINFAHL | -5.065 | 9.735 | 12.587 | - | + | - | + |

#### *REFERENCES CITED*


1. Daffern, N., Francino-Urdaniz, I. M., Baumer, Z. T. & Whitehead, T. A. Standardizing cassette-based deep mutagenesis by Golden Gate assembly. *Biotechnol. Bioeng.* **121**, 281–290 (2024).
2. Gibson, D. G. *et al.* Enzymatic assembly of DNA molecules up to several hundred kilobases. *Nat. Methods* **6**, 343–345 (2009).
3. Wrenbeck, E. E. *et al.* Plasmid-based one-pot saturation mutagenesis. *Nature Methods* vol. 13 928–930 Preprint at https://doi.org/10.1038/nmeth.4029 (2016).
4. Kirby, M. B., Medina-Cucurella, A. V., Baumer, Z. T. & Whitehead, T. A. Optimization of multi-site nicking mutagenesis for generation of large, user-defined combinatorial libraries. *Protein Eng. Des. Sel.* **34**, (2021).
5. Carrillo Rincón, A. F. & Farny, N. G. Unlocking the strength of inducible promoters in Gram-negative bacteria. *Microb. Biotechnol.* **16**, 961–976 (2023).
6. Cheetham, G. M. T., Jeruzalmi, D. & Steitz, T. A. Structural basis for initiation of transcription from an RNA polymerase–promoter complex. *Nature* **399**, 80–83 (1999).
7. Yin, Y. W. & Steitz, T. A. Structural basis for the transition from initiation to elongation transcription in T7 RNA polymerase. *Science* **298**, 1387–1395 (2002).
8. Tahirov, T. H. *et al.* Structure of a T7 RNA polymerase elongation complex at 2.9 Å resolution. *Nature* **420**, 43–50 (2002).
9. Durniak, K. J., Bailey, S. & Steitz, T. A. The structure of a transcribing T7 RNA polymerase in transition from initiation to elongation. *Science* **322**, 553–557 (2008).
10. Elmore, J. R., Furches, A., Wolff, G. N., Gorday, K. & Guss, A. M. Development of a high efficiency integration system and promoter library for rapid modification of Pseudomonas putida KT2440. *Metab Eng Commun* **5**, 1–8 (2017).
11. Choudhury, A. *et al.* CRISPR/Cas9 recombineering-mediated deep mutational scanning of essential genes in Escherichia coli. *Mol. Syst. Biol.* **16**, e9265 (2020).
12. Morgenthaler, A. B. *et al.* Mutations that improve efficiency of a weak-link enzyme are rare

compared to adaptive mutations elsewhere in the genome. *Elife* **8**, (2019).
